# The limits of evolutionary convergence in sympatry: reproductive interference and historical constraints leading to local diversity in warning traits

**DOI:** 10.1101/2021.01.22.427743

**Authors:** Ludovic Maisonneuve, Marianne Elias, Charline Smadi, Violaine Llaurens

## Abstract

Mutualistic interactions between defended species represent a striking case of evolutionary convergence in sympatry, driven by the increased protection against predators brought by mimicry in warning traits. However, such convergence is often limited: sympatric defended species frequently display different or imperfectly similar warning traits. The phylogenetic distance between sympatric species may indeed prevent evolution towards the exact same signal. Moreover, warning traits are also involved in mate recognition, so that trait convergence might result in heterospecific courtship and mating. Here, we develop a mathematical model to investigate the strength and direction of evolution of warning trait in defended species with different ancestral traits. Specifically, we determine the effect of phenotypic distances between ancestral trait states of sympatric defended species and of costs of heterospecific sexual interactions on imperfect mimicry and trait divergence. Our analytical results confirm that reproductive interference and historical constraints limit the convergence of warning traits, leading to either complete divergence, or imperfect mimicry. Our model reveals that imperfect mimicry evolves only when ancestral trait values differ between species due to historical constraints and highlights the importance of female and predator discriminations in the evolution of such imperfect mimicry. Our study thus provides new predictions on how reproductive interference interacts with historical constraints and may promote the emergence of novel warning traits, enhancing mimetic diversity.

## Introduction

Mutualistic interactions frequently drive convergent evolution of different traits in sympatric species. For example, avian vocal resemblance has been suggested to allow the formation of mixed-species flocks where all individuals from the different species may benefit from a reduced predation risk and an increase of their foraging (Goodale and Kotagama, 2006); likewise trait similarity between sympatric species may promote pollinator attraction in nectar-rewarding flowers (Benitez-Vieyra et al., 2007; Schemske, 1981). In aposematic species, warning traits are associated with defenses against predators, such as venom or disgusting taste. Because predators eventually learn such associations, convergence in warning trait among defended species facing similar communities of predators is often observed (Müllerian mimicry, see Sherratt (2008) for a review). Mimicry is certainly the best documented case of mutualistic interactions driving trait evolution in sympatric species and is observed in a wide range of organisms including plants (Lev-Yadun, 2009), mollusks (Cortesi and Cheney, 2010), vertebrates (Sanders et al., 2006; Springer and Smith-Vaniz, 1972) and insects (Mallet and Gilbert Jr., 1995). Field experiments report the intense selection exerted by predators favoring warning trait convergence in sympatry (Arias et al., 2016; Benson, 1972; Chouteau et al., 2016; Kapan, 2001; Mallet and Barton, 1989). Surprisingly, despite such intense selection, many sympatric defended species exhibit only imperfect resemblance (*e.g*. (Savage and Slowinski, 1992)) or even different warning traits (*e.g*. (Beccaloni, 1997)) (Briolat et al., 2019).

The level of trait convergence between sympatric species may vary depending on their level of phylogenetic relatedness. For instance, the significant phylogenetic signal observed on the warning trait of mimetic butterflies of the tribe Ithomiini (Chazot et al., 2014; Elias et al., 2008) suggests that historical constraints may limit the convergent evolution of warning traits. Such historical constraints are expected to be more different between distantly related species than closely-related ones, because closely-related species are expected to share similar ancestral trait values. These historical constraints may also imply differences between clades in both (1) the developmental pathway involved in the variation of warning traits or (2) the selective trade-offs between warning signals and other traits. For example, evolutionary history of different species influences their diet and since diet can influence the warning trait (Grill and Moore, 1998; Ojala et al., 2007), this may lead to different species-specific trade-offs limiting convergence between defended species. Historical constraints thus not only determine ancestral trait values, but also the evolvability of the traits in different species. Theoretical studies suggest that ancestral trait states may play a key role in the evolution of warning traits, because the convergence of trait can be facilitated by an initial resemblance between species (Balogh and Leimar, 2005; Franks and Noble, 2004; Franks and Sherratt, 2007). The initial resemblance between species, in the eyes of predators, depends on predator discrimination capacities, that then determines the strength of selection promoting convergence of warning traits. Other theoretical studies highlight that the level of standing genetic and phenotypic variance within species strongly influences convergence between species (Ruxton et al., 2008). The balance between (1) the shared predation pressure faced by individuals from different sympatric species and (2) the historical constraints within each species may thus strongly shape the level of evolutionary convergence in warning traits. This balance may also modify the direction of evolution of traits within the different defended species living in sympatry. While convergence (Sherratt, 2008) usually assumes a joint evolution of traits in several sympatric species toward resemblance (*e.g*. Flanagan et al. (2004); Symula et al. (2001)), resemblance might also emerge from advergence, whereby trait evolution occurs in a given species (*i.e*. the ‘mimic’ species), leading to high similarity to the ancestral trait displayed in another species (i.e. the ‘model’ species) (see (Dalziell and Welbergen, 2016) for the terminology).

Moreover, the convergence of warning traits in different species may entail costs due to behavioral interference, thereby limiting positive selection on trait resemblance. Warning traits are indeed frequently involved in species recognition (Jiggins et al., 2001; Kronforst et al., 2006; Merrill et al., 2014; Naisbit et al., 2001), leading to increased risk of confusion in mimetic species during sexual interactions. Such risk might be even higher between closely-related species, which are more likely to share multiple similar traits because of common ancestry. Species sharing similar warning traits may thus be exposed to substantial reproductive interference incurring fitness costs during mate acquisition due to interspecific interactions, including heterospecific courtship and mating as well as heterospecific male rivalry (Gröning and Hochkirch, 2008). Empirical examples of such reproductive interferences in Müllerian mimetic systems have been reported in the literature (Estrada and Jiggins, 2008; Vasconcellos-Neto and Brown, 1982). However, empirical studies precisely estimating the level of reproductive interference in sympatric species are scarce. Pheromone differences between mimetic species have been documented to limit the rate of erroneous mating (see Darragh et al. (2017); González-Rojas et al. (2020) for empirical examples in *Heliconius* butterflies). However, the pheromones of day-flying butterflies usually act as short-distance cues that may be perceived only during courtship (Mérot et al., 2015). Females deceived by the color pattern of the heterospecific males may have already spent time and energy or may need to deploy substantial efforts to avoid heterospecific mating. Therefore, females may still suffer from costs associated to reproductive interference, even if females refuse mating with heterospecific males. When females are courted by heterospecific males displaying their preferred cue before being rejected, this also results in increased costs associated with mate searching in males (i.e. signal jamming in (Gröning and Hochkirch, 2008)).

Reproductive interference can generate reproductive character displacement (Gröning and Hochkirch, 2008; Kyogoku, 2015), whereby reproductive traits are more dissimilar between species in sympatric than in allopatric populations (Brown and Wilson, 1956). Such reproductive character displacement may thus impair convergence driven by mutualistic interactions. Theoretical studies have investigated how the evolution of female preferences may promote reproductive character displacement in males (McPeek and Gavrilets, 2006; Yamaguchi and Iwasa, 2013): reproductive interference costs are predicted to favor divergence between female preference and trait displayed by heterospecifics, because this reduces mating attempts with heterospecifics, and therefore promotes the divergence of reproductive traits between conspecific and heterospecific males through sexual selection. Female discrimination then determines the level of divergence between female preference and trait displayed by heterospecifics necessary to limit the cost of reproductive interference (McPeek and Gavrilets, 2006; Yamaguchi and Iwasa, 2013). Numerical simulations assuming two discrete warning traits and fixed warning trait-based assortative mating show that reproductive interference may impair the convergence of warning traits (Boussens-Dumon and Llaurens, 2021). Nevertheless, understanding the impact of reproductive interference on the evolution of warning trait requires to specifically explore the evolution of female preference towards this trait. Moreover, the outcomes of these antagonistic selective forces might range from trait divergence to full convergence, through limited convergence and cannot be investigated in models assuming only discrete and well-differentiated warning traits, calling for a theoretical framework providing general expectations on the gradual evolution of convergent traits.

Here, we thus investigate the selective pressure limiting the convergence of traits involved in mimetic interactions, by building a mathematical model that describes the evolution of quantitative traits in two sympatric species engaged in mimetic interaction. We specifically study the evolution of (1) the quantitative trait *t* involved in mimetic interaction, displayed in both males and females and (2) the preference *p*, which value indicates the male trait value preferred by the female. We assume that individuals from different species gain protection from predators, by sharing similar warning trait values with other defended individuals living in the same environment, whatever species they belong to. However, trait similarity between species generates fitness costs for females *via* reproductive interference (McPeek and Gavrilets, 2006; Yamaguchi and Iwasa, 2013). We neglect fitness costs of reproductive interference acting on males (McPeek and Gavrilets, 2006; Yamaguchi and Iwasa, 2013), reflecting the asymmetrical investment in reproduction between sexes observed in numerous species (Trivers, 2017). We assume that a parameter *c_RI_* modulates the strength of reproductive interference. The strength of reproductive interference depends on the degree of similarity between species. Because the selective forces acting on warning traits strongly depend on the sensitivity of both females and predators, we test the effect of their discrimination capacity on convergent evolution. We then investigate the interactions between these opposed selective forces with the effect of historical constraints, reflecting evolutionary history, by assuming different ancestral trait values in the two interacting species, as well as stabilizing selection promoting these ancestral values within each species. Using weak selection approximation (Barton and Turelli, 1991; Kirkpatrick et al., 2002), we obtain equations describing the evolution of the mean trait and mean preference values in both species. We then use analytical results and numerical analyses to investigate the effect of reproductive interference on the convergence of trait, depending on different ecological factors.

## Methods

We consider two sympatric species, called species 1 and 2. In species *i* for *i* ∈ {1, 2}, males and females display a warning trait *t_i_*. We assume that only females express a mating preference for males. The value of female preference *p_i_* indicates the value in male trait triggering the highest attraction of the female. We investigate the evolution of the warning trait and preference within each species, influenced by both natural selection and mate choice.

### Model structure

We assume constant population size in the two species and balanced sex ratio. We consider discrete and non-overlapping generations. The offspring in each new generation are produced by sexual reproduction between males and females from the previous generation, following a Wright-Fisher Model (Fisher, 1930; Wright, 1931). Following the framework from Barton and Turelli (1991); Kirkpatrick et al. (2002), in species *i* for *i* ∈ {1, 2}, we assume that the distribution of traits in the offspring depend on the so-called group absolute fitness *W^i^*(*t_m_, t_f_, p_f_*) of the parental generation. This absolute group fitness accounts for the trait values displayed by the males (*t_m_*) and the females (*t_f_*), as well as on the preference of the females (*p_f_*) producing the offspring generation. This group absolute fitness Wi thus describes the effect of selection acting on viability and fecundity, as well as the sexual selection due to mate preference, on the evolution of trait and preference in the population, as detailed below.

### Life cycle

Because warning traits are involved in survival and mate choice, the group absolute fitness is a function of male and female traits and female preference. Following Pomiankowski and Iwasa (1993) (see Equation A1), the group absolute fitness associated with each pair is assumed to be given by the product of different fitness terms and of one term describing mate preference. For *i* ∈ {1, 2}, the group absolute fitness associated with a pair consisting of a male with trait *t_m_* and a female with trait *t_f_* and preference *p_f_* is assumed to be given by:

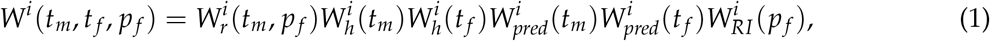

where 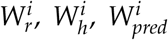 and 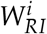 describes respectively the action of mate preference, historical constraints, predation and reproduction.

#### Mate preference

In each species *i* ∈ {1, 2}, the contribution to the next generation of a mating between a male with trait *t_i_* and a female with preference *p_i_* due to mate preference is assumed to be given by

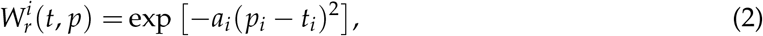

where female discrimination *a_i_* > 0, assumed constant among conspecific females, quantifies how much females of species *i* reject males with a non-preferred trait value.

#### Ancestral trait value

Phenotypic evolution in both species away from their ancestral trait is limited by historical constraints, specific to each species. The phenotypic evolution thus strongly depends on ancestral trait values in both species *t*_*a*1_, *t*_*a*2_, as well as on the stabilizing selection promoting this ancestral trait value *t_ai_* within each species. The strength of the stabilizing selection within each species *i* depends on the coefficient *s_i_*. The fitness component due to historical constrainsts is thus assumed to be given by:

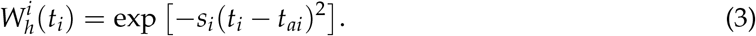

#### Predation depending on the level of mimicry of trait t

Within each species, the evolution of the trait *t*, expressed by males and females from a species, is strongly influenced by the trait displayed in the other species. Müllerian mimicry indeed generates positive density-dependent selection (Benson, 1972; Chouteau et al., 2016; Mallet and Barton, 1989), due to predator learning. This density-dependence is non linear and is often modeled as an hyperbolic decrease in mortality (see (Joron and Iwasa, 2005; Llaurens et al., 2013) for example). The impact of predation on the fitness of an individual displaying the trait value *t* is assumed to be given by:

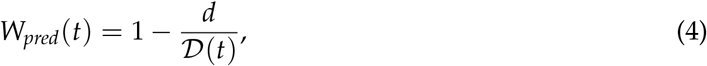

where *d* ∈ (0, 1) the basal predation rate, and 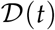 is the level of protection of an individual with trait *t* increasing with the density of resembling individuals:

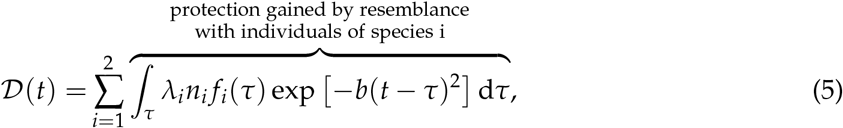

where for each *i* ∈ {1, 2}, *f_i_* is the distribution of traits, and *n*_1_ is the density of individuals, in species *i*. The density-dependence is modulated by the individual levels of defense *λ*_1_ and *λ*_2_, assumed constant among individuals of species 1 and 2, respectively, shaping predator deterrence: the higher the defense, the quicker predators learn. The protection gained against predators then depends on the level of resemblance among defended prey, as perceived by predators, and on the number of individuals sharing similar trait values. Due to the positive density-dependent selection, *λ_i_n_i_* is then the population defense level in species *i*. exp [–*b*(*t* – *τ*)^2^] describes how much predators perceive the trait values *t* and *τ* as similar. The predator discrimination coefficient *b* thus quantifies how much predators discriminate different trait values.

#### Cost induced by reproductive interference

Because heterospecific males may resemble conspecific males, females will suffer from reproductive interference generated by erroneous mating attempts with heterospecific males (see (Gröning and Hochkirch, 2008) for a review of reproductive interference costs). The risk of heterospecific mating depends on the relative densities of heterospecific and conspecific males. We assume a balanced sex-ratio within each species *i.e*. the density of males in species *i* is *n_i_*/2, for *i* ∈ {1, 2}. However, we also consider the capacity of females to recognize conspecific males using alternative cues (pheromones for example). In the model, the investment of females in interspecific mating interaction is captured by the strength of reproductive interference *c_RI_* ∈ [0, 1]. This cost of reproductive interference incurred to females can be reduced when female choice is also based on alternative cues differing between mimetic species. Using Equation (1b) in (Yamaguchi and Iwasa, 2013) the fitness of a female of species *i* ∈ {1, 2} with preference *p_i_* is modulated by:

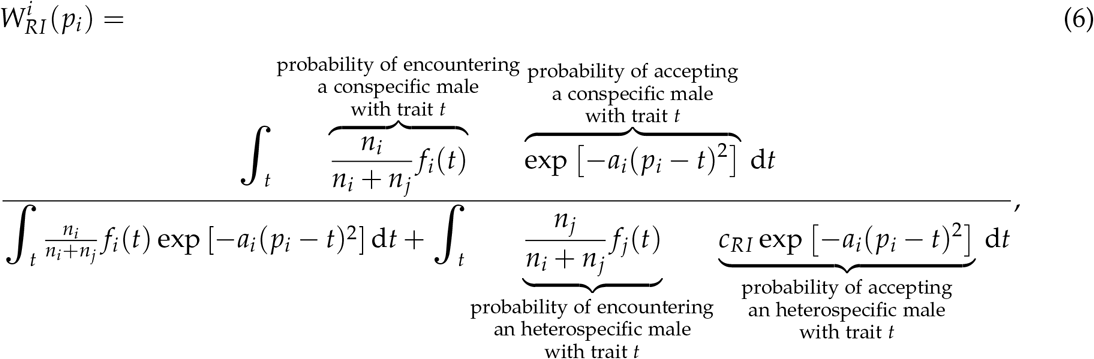

with *j* ∈ {1, 2} with *j* ≠ *i*.

### Approximation of the evolutionary dynamics

We assume that in each species the trait and preference are quantitative traits, with an autosomal polygenic basis, and additive effects. We assume that male and female traits have the same genetic basis.

We assume weak natural and sexual selective pressures (Iwasa et al., 1991; Pomiankowski and Iwasa, 1993) implying that the variance of trait and preference is small relative to the curvature of the fitness function in each species (see Appendix 1). Using the Price’s theorem (see Rice (2004) for instance), we can approximate the change in the mean values of traits 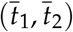 and preferences 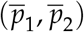 in both species, after the natural and sexual selection respectively, by:

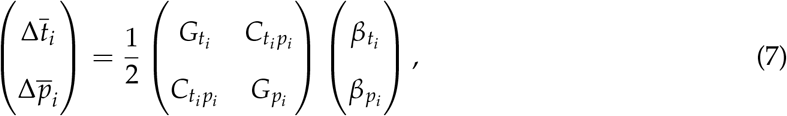

where for *i* ∈ {1, 2} *G_t_i__* and *G_p_i__* are the additive genetic variances of *t_i_* and *p_i_* and *C_t_i_p_i__* is the additive genetic covariance between *t_i_* and *p_i_*. *β_t_i__* and *β_p_i__* describe the selective forces acting on the trait *t_i_* and the preference *p_i_* respectively and are given by:

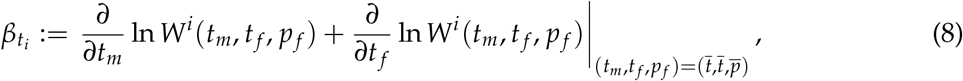

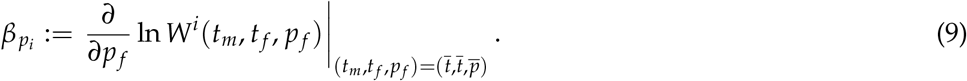

Under a weak selection hypothesis, genetic correlations generated by selection and nonrandom mating quickly reach equilibrium (Nagylaki, 1993) and can thus be approximated by their equilibrium values.

Following (Iwasa et al., 1991), we assume that for each *i* ∈ {1, 2}, *G_t_i__* and *G_p_i__* are positive constants maintained by an equilibrium between selection and recurrent mutations. Under weak selection, for each *i* ∈ {1, 2}, the genetic covariance between *t_i_* and *p_i_* can be approximated by (see Appendix 4):

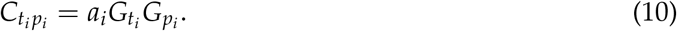

When the variance of trait and preference is small relative to the curvature of the fitness function, for each *i* ∈ {1, 2}, *C_t_i_p_i__* is small in comparison with *G_t_i__* and *G_p_i__*, allowing us to approximate the change in the mean values of trait and preference in each species *i* ∈ {1, 2} by:

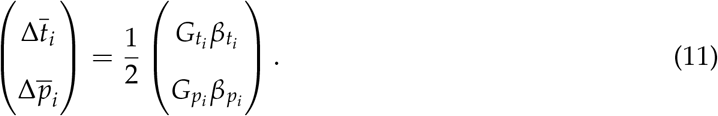

Using Equation (11) we derive 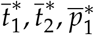 and 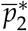 the mean traits and preferences at equilibrium (see Appendix 5 & 6).

All variables and parameters used in the model are summed up in Table 1. The effect of different parameters on the evolutionary outcome are presented in Appendix 5.4 and 6.3.

**Table 1:** Description of variables and parameters used in the model. The subscript *i* ∈ {1, 2} denotes the identity of the species.

| Abbreviation | Description |
| --- | --- |
| $\bar{t}_i/\bar{p}_i$ | Mean trait/preference value in species $i$ |
| $G_{t_i}/G_{p_i}$ | Genetic variance of trait $t_i$/preference $p_i$ |
| $C_{t_i p_i}$ | Genetic covariance between trait $t_i$ and preference $p_i$ |
| $\beta_{t_i}/\beta_{p_i}$ | Selection coefficient on trait $t_i$/preference $p_i$ |
| $a_i$ | Female discrimination in species $i$ |
| $s_i$ | Strength of stabilizing selection due to historical constraints on trait $t_i$ |
| $t_{ai}$ | Ancestral trait in species $i$ |
| $d$ | Basal predation rate |
| $b$ | Predator discrimination |
| $n_i$ | Density of species $i$ |
| $\lambda_i$ | Individual defense level in species $i$ |
| $\lambda_i n_i$ | Population defense level in species $i$ |
| $c_{RI}$ | Strength of reproductive interference |

## Results

### Evolution of the warning trait in a mimetic species in sympatry with a model species

To identify the general impact of the strength of reproductive interference *c_RI_* on the phenotypic distance between the two species 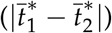, we first look at the analytical resolution assuming that trait and preference are fixed in species 2 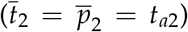 and weak female and predator discriminations (*a*_1_ = *O*(*ε*) and *b* = *O*(*ε*)) (see Appendix 5.2). This cover cases where species 2 is well defended and more abundant than species 1 (*n*_2_ ≫ *n*_1_) as in classical *mimic/model* interactions between species.

#### Analytical expression of the mean trait and preference values

Assuming weak female and predator discriminations, the mean trait and preference values both converge to the equilibrium values 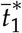 and 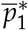 with

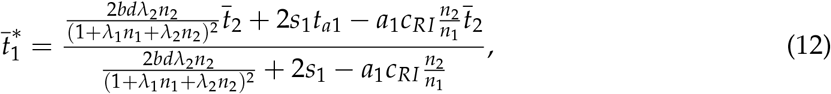

and

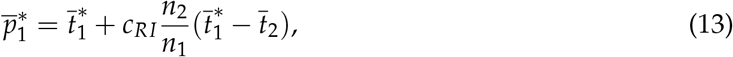

when

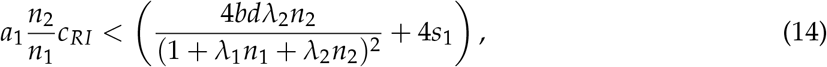

see Appendix 5.2.

These analytical expressions allow us to predict on the level of resemblance between the trait displayed in species 1 and the fixed trait exhibited in the *model* species 2 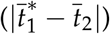 and to study the impact of the different evolutionary forces on the advergence between *mimic* and *model*.

However, when (14) is not verified, the distances between mean trait and preference values in species 1 and 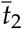 become very large (not of order 1), and mimicry does not emerge (see Appendix 5.2).

#### Reproductive interference limits mimicry

Selection exerted by predators favors the advergence of trait in species 1 toward the fixed trait value exhibited in the *model* species 2. When (14) is verified, the level of advergence toward 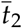 is given by:

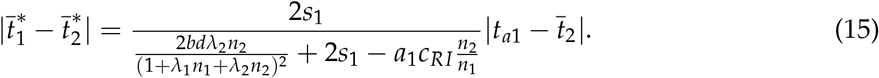

Hence, if we assume no reproductive interference (*c_RI_* = 0), we have 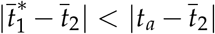, implying that the trait displayed in species 1 gets closer to the trait displayed in species 2. We then observe different evolutionary outcomes ranging from (a) *mimicry* to (b) *imperfect mimicry*, see Figures 1(a) and 1(b). Mimicry in species 1 becomes nearly perfect 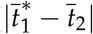 close to zero) when the strength of selection due to predation (2*bdλ*_2_*n*_2_/(1 + *λ*_1_*n*_1_ + *λ*_2_*n*_2_)^2^) is large enough, as compared to the historical constraints limiting the evolution of the trait in species 1 (*s*_1_) (outcome (a) *mimicry* see Figure 1(a)).

**Figure 1:**
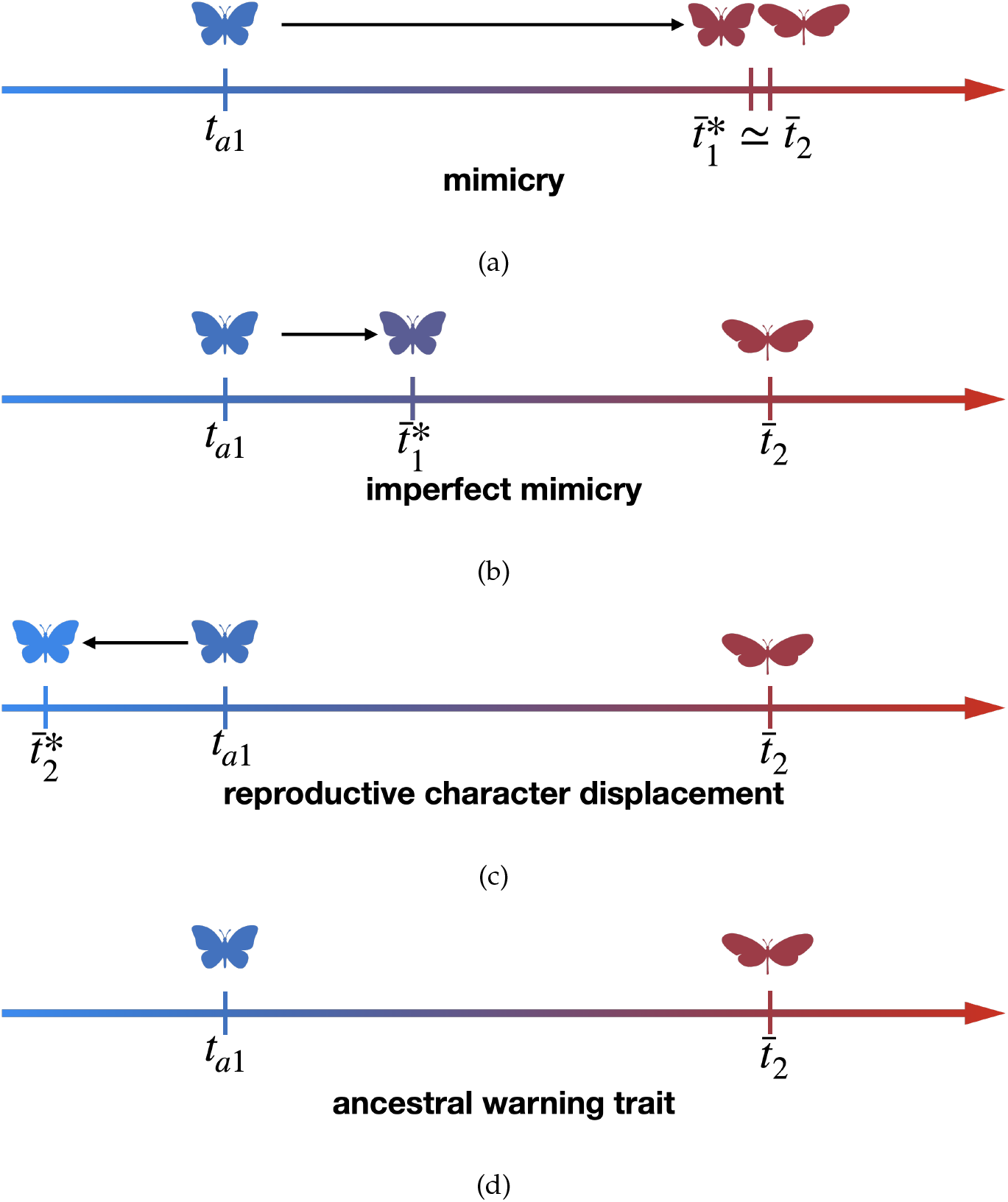
Illustration of four approximate patterns referred in this paper: (a) *mimicry:* the value of the trait in species 1 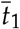 becomes very close to the mean value displayed in species 2 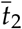, (b) *imperfect mimicry:* the value of the trait in species 1 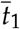 gets closer but stays distant from the mean value displayed in species 2 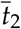, (c) *warning trait displacement:* the value of the trait species 1 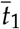 diverges away from the mean value displayed in species 2 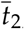, (d) *ancestral warning trait:* the value of the trait in species 1 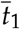 stays very close to the ancestral trait value *t*_*a*1_.

However, assuming reproductive interference between females from species 1 and males from species 2 impairs advergence. When reproductive interference is non null but has a limited strength, satisfying Inequality (14), Equation (15) implies that 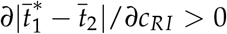 (see Appendix 5.4.1). Reproductive interference thus increases the distance between the traits displayed in both species, leading to imperfect mimicry in species 1. Reproductive interference promotes the evolution of preference in the opposite direction of the trait displayed by heterospecific males (13). Because female preference generates sexual selection on male traits, reproductive interference promotes phenotypic divergence between both species (see Equation (13)). Thus reproductive interference limits Müllerian mimicry.

However, when the cost associated with reproductive interference crosses a threshold and (14) is verified, *i.e*. when

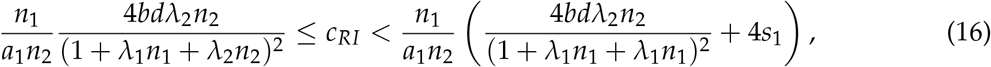

then

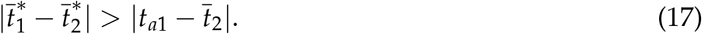

When assuming such an elevated cost of reproductive interference, imperfect mimicry is thus no longer observed, and reproductive interference rather promotes warning trait displacement. The trait in species 1 diverges away from the trait displayed in species 2 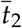 (see Figure 1(c) for an illustration).

When inequality (14) is not satisfied, the phenotypic distance between both species becomes very large. This very large divergence is biologically unrealistic but suggests that reproductive interference can promote phenotypic divergence between defended species living in sympatry. This unrealistic divergence stems from the weak female discrimination (*a*_1_ = *O*(*ε*)) assumed: since females have low discrimination (because *a*_1_ is low), females almost always accept heterospecific males, except when the difference between female preference in species 1 and the trait displayed in species 2 is very high. Reproductive interference promotes female preference that limits fitness costs due to reproductive interference, and therefore promotes a large distance between females preference value in species 1 and the value of the trait displayed in species 2. Relaxing the weak female and predator discriminations hypothesis, *i.e*. assuming that *a*_1_ = *O*(1) and *b* = *O*(1), confirms that reproductive interference limits mimicry in species 1 (see Figure 2). However, in this case, when a strong divergence is favored, this divergence becomes high but stays of order *O*(1). Indeed, as female discrimination is high, this divergence strongly reduces fitness cost due to reproductive interference. Therefore, stabilizing historical constraints on the trait becomes more important than reproductive interference, thereby preventing very large divergence. Figure 2 shows that numerical simulations with parameter values matching weak female and predator discriminations provide similar predictions as the analytical approximation obtained under the same hypotheses.

**Figure 2:**
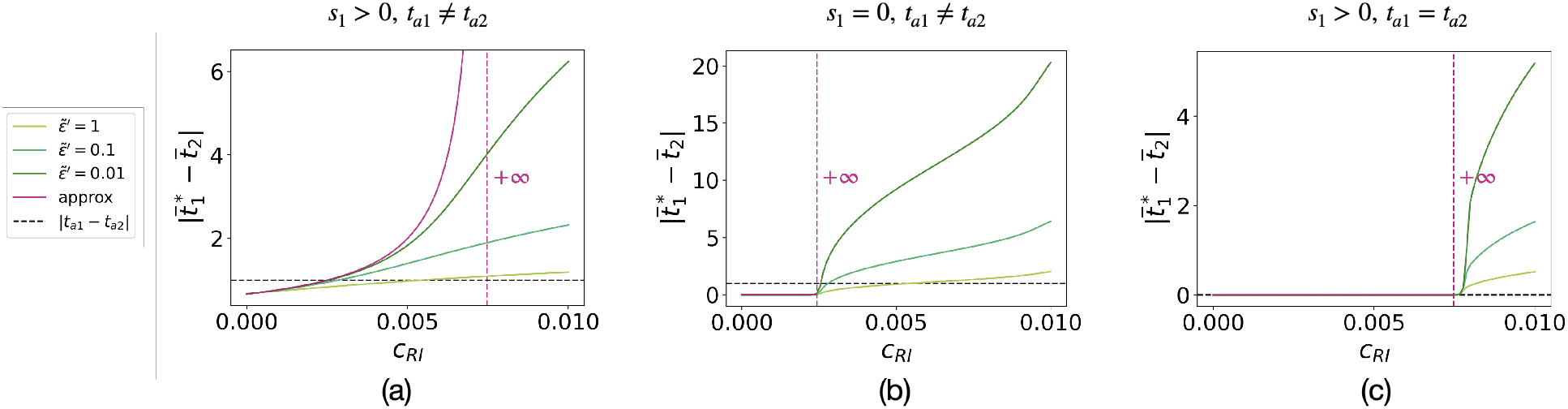
Influence of the strength of reproductive interference *c_RI_* on the phenotypic distances between the two species 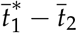 when trait in species 2 is fixed 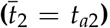, using the analytical approximation (purple curve) or numerical simulations (green curves). The different values of 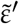 allows to investigate the intermediate case between weak and strong female and predator discriminations. We assume (a) 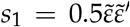, *t*_*a*1_ = 0, *t*_*a*2_ = 1, (b) *s*_1_ = 0, *t*_*a*1_ = 0, *t*_*a*2_ = 1, and (c) 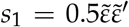, *t*_*a*1_ = *t*_*a*2_ = 1 with 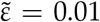. We also assume: *G*_*t*_1__ = *G*_*p*_1__ = 0.01, 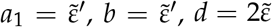, *λ*_1_ = 0.1, *λ*_2_ = 0.1, *n*_1_ = 10, *n*_2_ = 20. Analytical approximation curves are obtained with 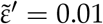.

#### Historical constraints allow for the evolution of imperfect mimicry

Our previous results highlight that reproductive interference limits the convergence of warning traits. However, the effect of reproductive interference on trait divergence strongly depends on the historical constraints (*s*_1_), generating a stabilizing selection promoting the ancestral trait value (*t*_*a*1_). Strong historical constraints promote the display of the ancestral trait in both species (see Equation 15 and Figure 1(d) for an illustration when species 2 is fixed). The effect of historical constraints on the level of trait divergence depends then on the distance between the ancestral trait values in species 1 and 2. When predator pressure exceeds reproductive interference, historical constraints limit the convergence of trait between both species ancestrally displaying different traits (see Appendix 5.4.4). By contrast, when reproductive interference exceeds predator pressure and promotes warning trait displacement, historical constraints may limit the divergence of trait between both species (see Appendix 5.4.4). Assuming historical constraints (*s*_1_ > 0) and when the ancestral trait values of the two species differ (*t*_*a*1_ ≠ *t*_*a*2_), an increase in strength of reproductive interference leads to a progressive increase in the phenotypic distance between both species until the phenotypic distance between both species becomes very large (see purple curve in Figure 2(a)).

Surprisingly, in the absence of historical constraints (*s*_1_ = 0) or when the ancestral trait values are the same in both species (*t*_*a*1_ = *t*_*a*2_), 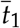 is either equal to 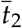 when (14) is verified, or is very large, when (14) is not verified (see purple curve in Figures 2(b) and 2(c)). Therefore an increase in the strength of reproductive interference (*c_RI_*) has no effect on the phenotypic distance between both species, as long as this strength remains below a threshold. This effect is also observed assuming strong female and predator discriminations (see green curves in Figures 2(b) and 2(c)).

However, when the strength of reproductive interference (*c_RI_*) is greater than this threshold, assuming weak female and predator discriminations, the phenotypic distance between both species becomes instantaneously very large. A similar trend is observed when female and predator discriminations are strong: the phenotypic distance is null when the strength of reproductive interference remains below a threshold, but it quickly increases to a high value when the strength of reproductive interference crosses the threshold (see green curves in Figures 2(b) and 2(c)).

Our results highlight that historical constraints promoting ancestral traits strongly modulate the effect of reproductive interference on the convergence of warning traits. Surprisingly, drastic divergence might be promoted by a strong strength of reproductive interference, even when the ancestral phenotypes are the same in the two interacting species.

Overall, our analytical results reveal the mechanisms underlying trait and preference evolution. However, these analytical results are obtained under restrictive hypotheses: we assumed fixed trait and preference in species 2 and weak female and predator discrimination. To relax those hypotheses, we then study the joint evolution of traits and preferences in both species in the following sections. We verified that all results obtained in previous sections are maintained when traits and preferences jointly evolve in the two sympatric species (see Appendix 6.3 and 6.4)..

### Joint evolution of mimicry between two interacting species

In this section, we now focus on the general case where traits and preferences co-evolve in both species.

#### Higher discrimination in female than in predator does not always favor the convergence of warning traits between two interacting species

The joint evolution of the traits in both species is shaped by two antagonistic evolutionary forces, generated by reproductive interference and Müllerian mimicry, respectively. Reproductive interference indirectly limits mimicry by impacting females’ preference. Therefore, female discrimination *a*_1_ and *a*_2_ may be a key feature to understand the evolution of the trait within each species. The selection exerted by predation also depends on predator discrimination. Assuming a fixed level of reproductive interference (*c_RI_* = 0.002), we thus investigate the impact of the strength of female discrimination, assumed equal in both species *a*_1_ = *a*_2_ = *a*, and of predator discrimination coefficient *b* on the evolution of the warning trait.

When female and predator discriminations are low (*a* and *b* approximately lower than 3), higher predator than female discrimination favors the convergence of warning traits. Indeed, when female and predator discriminations are low, selection due to predation and reproductive interference is limited and increases with female and predator discriminations, respectively. Females are not discriminant (*a* is low) and tend to accept all encountered males, including heterospecific males, whatever the direction of their preference. The difference in fitness cost due to reproductive interference between females with different preferences is then low, leading to low divergent selection generated by reproductive interference. With a higher level of female discrimination, fitness cost due to reproductive interference depends more on the direction of preference, leading to higher selection caused by reproductive interference. A similar reasoning on the difference in fitness cost due to predation between individuals displaying different traits explain that selection promoting convergence, due to predation, increases with the strength of predator discrimination. Therefore, higher predator than female discrimination entails higher selection due to predation than selection due to reproductive interference and promotes mimicry.

By contrast, with higher female discrimination (*a* approximately greater than 3) and lower predator discrimination (*b* approximately lower than 3), mimicry becomes more likely (Figure 3). Higher levels of female discrimination allow females to accurately distinguish between conspecific and heterospecific males even when they display similar traits. Accurate choice by females allows both species to harbor similar traits from the point of view of predators, without entailing heterospecific mating, relaxing divergent selection generated by reproductive interference.

**Figure 3:**
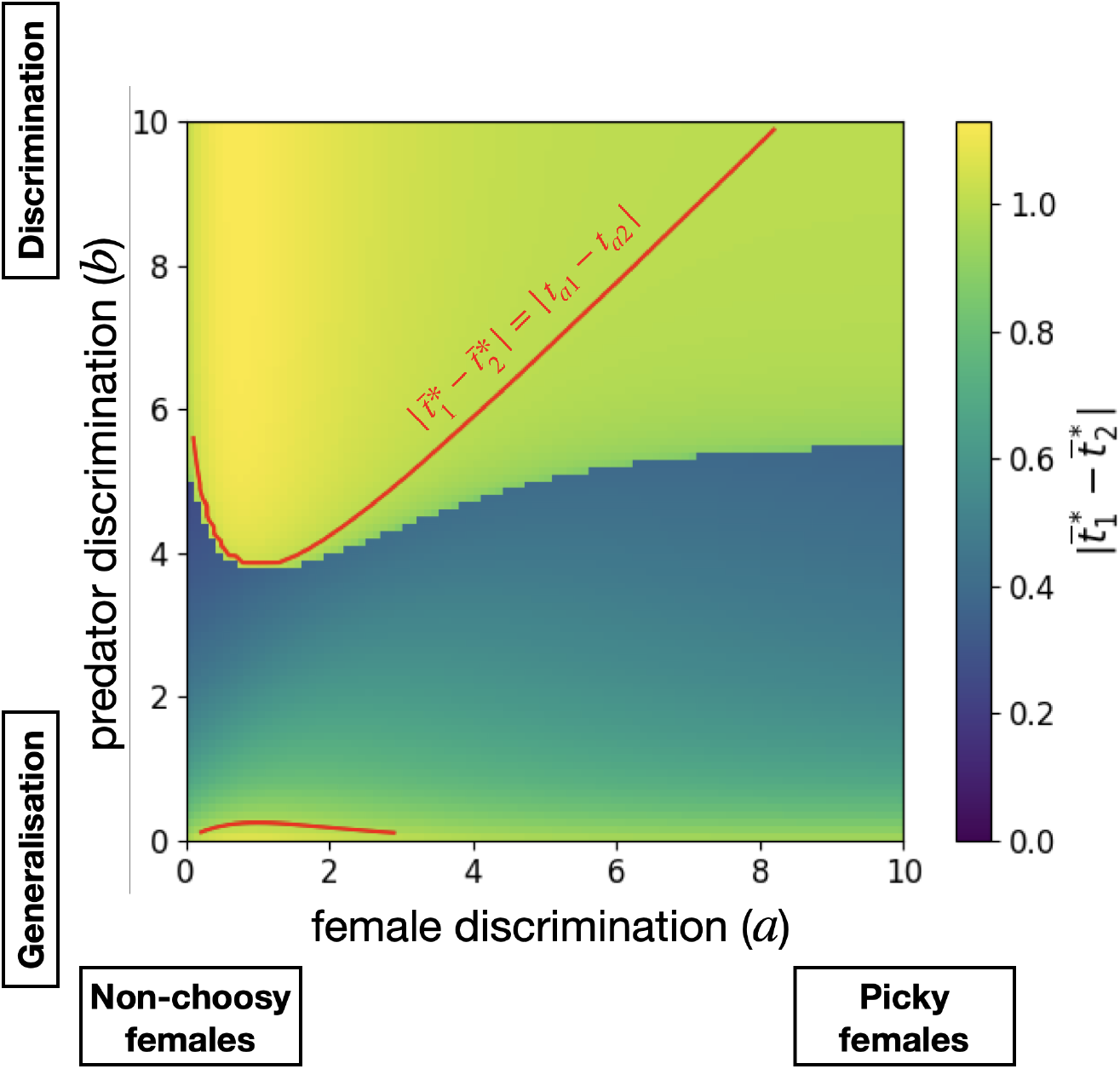
Influence of female and predator discriminations (*a*_1_ = *a*_2_ = *a* and *b*) on the phenotypic distance between the two species 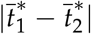. The red solid line shows the case where the phenotypic distance between the two species is equal to the ancestral phenotypic distance 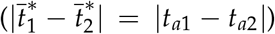. We assume: *G*_*t*_1__ = *G*_*p*_1__ = *G*_*t*_2__ = *G*_*p*_2__ = 0.01, *c_RI_* = 0.002, *d* = 0.02, *λ*_1_ = *λ*_2_ = 0.1, *n*_1_ = *n*_2_ = 20, *s*_1_ = *s*_2_ = 0.005, *t*_*a*1_ = 0, *t*_*a*2_ = 1.

Surprisingly, when predator discrimination increases above a certain threshold, increased discrimination no longer promotes accurate mimicry (see sharp transition in Figure 3). When *b* is approximately greater than 5.5, mimicry is limited, even without reproductive interference (*c_RI_* = 0), because of historical constraints (see Figure A21). For intermediate predator discrimination (*b* ≈ 5), mimicry is limited when reproductive interference is strong and makes similarity too costly for females (*a* ≈ 1) (Figure 3).

When reproductive interference limits mimicry, it generally leads to warning trait displacement 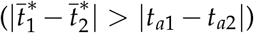, when female discrimination is low. Under low female discrimination, reproductive interference promotes a large distance between female preference value in species 1 and the value of the trait displayed in species 2, therefore increasing phenotypic distance between the two species.

#### Reproductive interference can modify the model/mimic relationship

The population defense levels in both species, *i.e. λ_i_n_i_, i* ∈ {1, 2}, are likely to impact the joint evolution of traits in both species. To investigate how the relative population defense levels of the two species affect the joint evolution of traits, we studied the phenotype at equilibrium in both species and also the phenotypic distance between the two species, for different values of the two components of the population defense level in species 1: the individual defense level (*λ*_1_) and the density (*n*_1_). Here we assumed that species 2 is already well protected (*λ*_2_ = 0.1, *n*_2_ = 10).

*When assuming no reproductive interference* (*c_RI_* = 0), the trait of the less defended species adverges towards the ancestral trait of the most defended species. In Figures 4(a) and 4(b), individuals from the poorly defended species 1 (i.e. when *λ*_1_*n*_1_ is low) get weak protection from conspecific individuals and thus have a greater advantage to look similar to individuals of species 2. Convergence of warning traits is thus more likely to happen when species 1 is weakly defended (*λ*_1_*n*_1_ small) (see Figure 4(c)). The more species 1 is defended, *i.e*. the greater *λ*_1_*n*_1_ is, the closer its mean trait value is to the ancestral trait value *t*_*a*1_ (see Figure 4(a)). Such increase in the defense level of species 1 also impacts the evolution of trait in the sympatric species 2 (see Figure 4(b)): when the individual defense level in species 1 (*λ*_1_) is below a threshold, the more individuals from species 1 are protected, the more the mean trait value in species 2 moves away from its ancestral trait (*t*_*a*2_). Surprisingly, above this threshold, the better protected species 1 is, the closer the mean trait value in species 2 gets to its ancestral trait value (*t*_*a*2_). As the mean trait value in species 1 becomes very close to the ancestral trait value *t*_*a*1_, trait values in species 2 leading to protection from heterospecific matings necessitate a great departure from the ancestral trait *t*_*a*2_. Nevertheless, historical constraints still prevent the trait in species 2 to evolve too far away from its ancestral trait value *t*_*a*2_.

**Figure 4:**
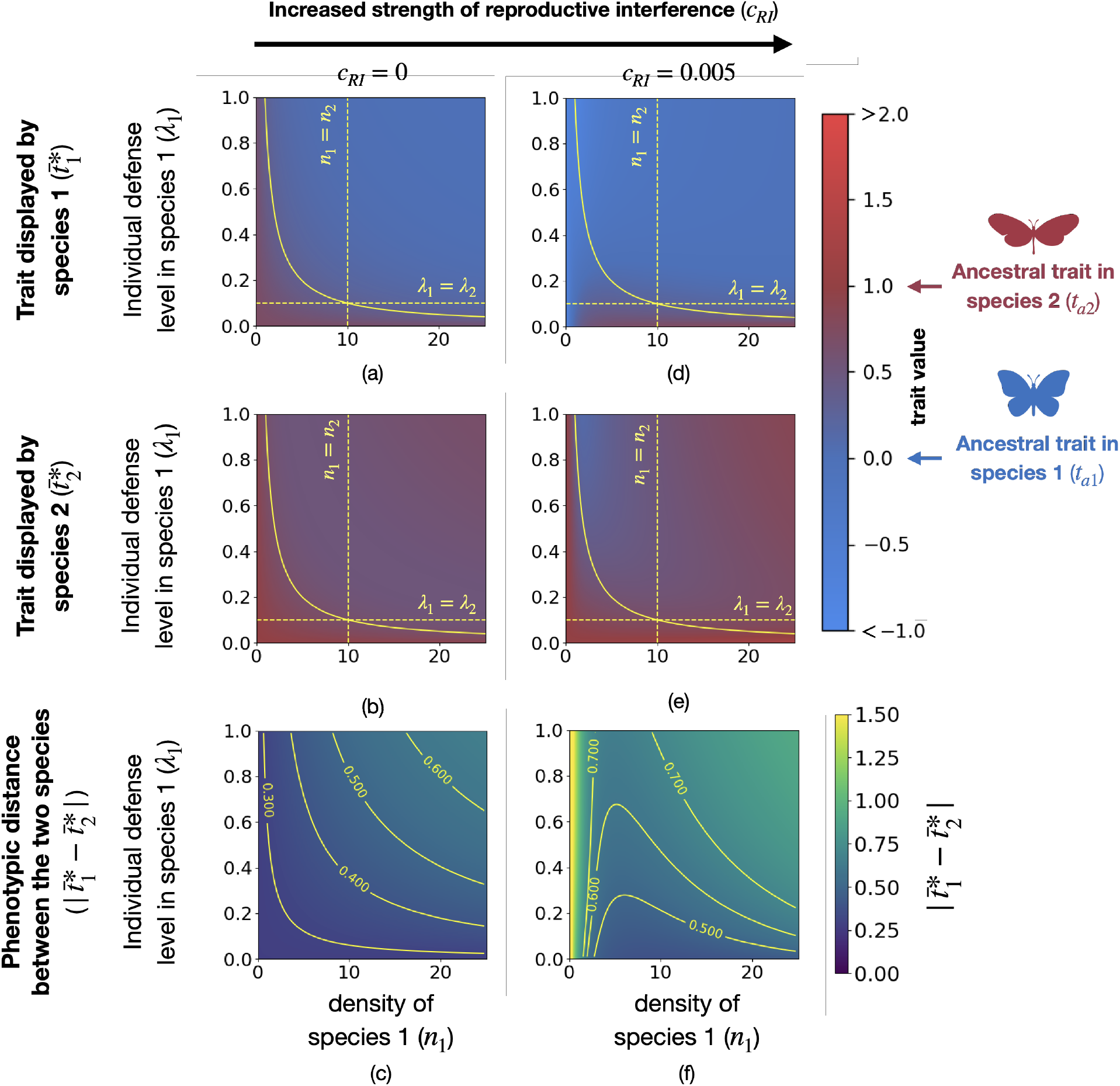
Influence of the density and of the individual defense level in species 1 (*n*_1_ and *λ*_1_) on the traits displayed in both species and on the phenotypic distance between the two species 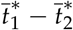, for different strengths of reproductive interference (*c_RI_*). (a)(b)(d)(e) Trait values greater than 2 (resp. lower than −1) are shown in red (resp. blue). The yellow solid line shows the case where both species have the same level of defense (*λ*_1_*n*_1_ = *λ*_2_*n*_2_). Below (resp. above) this line species 1 has a lower (resp. higher) level of defense than species 2. (c)(f) Phenotypic distances greater than 1.5 are shown in yellow. Yellow lines indicate equal levels of 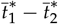. Different values of strengths of reproductive interference are assumed: (a), (b) and (c) *c_RI_* = 0, (d), (e) and (f) *c_RI_* = 0.005. We assume: *G*_*t*_1__ = *G*_*p*_1__ = *G*_*t*_2__ = *G*_*p*_2__ = 0.01, *a*_1_ = *a*_2_ = 1, *d* = 0.05, *b* = 1, *λ*_2_ = 0.1, *n*_2_ = 10, *s*_1_ = *s*_2_ = 0.005, *t*_*a*1_ = 0, *t*_*a*2_ = 1.

*When assuming positive strength of reproductive interference* (*c_RI_* > 0), advergence in species 1 toward the trait displayed in species 2 is observed when the individual defense level in species 1 is low (*λ*_1_ approximately lower than 0.1) and when the density in species 1 is sufficiently large (*n*_1_ approximately greater than 2). In this case, the population defense level in species 1 (*λ*_1_*n*_1_) is low, the protection gained by positive frequency-dependent selection within species is low, and the advergence toward species 2 is thus strongly promoted. Surprisingly, advergence is impaired for similar values of defense level, when the density of species 1 is low (*n*_1_ approximately lower than 2). When the density of species 1 is low, females pay higher fitness costs due to reproductive interference, because they encounter more often heterospecific than conspecific males. Altogether, our results suggest that advergence of the warning traits is likely to happen for low level of individual defense in species 1 (i.e. Batesian (*λ*_1_ = 0) or quasi-Batesian (*λ*_1_ > 0 but small) mimicry) and when the density of species 1 is high.

The trait value of species 1 does not always converge toward the trait value initially displayed in species 2 (*t*_*a*2_). On the contrary, individuals of species 2 can mimic individuals of species 1 (see blue zone in Figure 4(e)), when the defense level of individuals of species 1 is high and when species 1 is rare. Because individuals from both species are well defended (high *λ*_1_ and *λ*_2_), individuals of both species benefit from looking similar. However, when species 1 is rarer, this leads to an increased cost of reproductive interference in species 1, inhibiting convergence towards the ancestral trait value displayed in the alternative species (*t*_*a*2_) (see Figure 4(d)). Since predation pressure promotes convergence of traits of both species, the mean trait in species 2 becomes closer to species 1 ancestral trait value *t*_*a*1_. Surprisingly assuming weak female and predator discriminations, such advergence also happens when individuals of species 2 are more defended than individuals from species 1, *i.e. λ*_2_*n*_2_ > *λ*_1_*n*_1_ (see blue zones in Figures A22 (c) and (d) below the yellow solid line). By contrast, when the density in species 1 exceeds the density in species 2, individuals from both species exhibit traits close to their ancestral traits. Both species are well-protected and then gain little from mimicry. Because unbalanced relative density ratio leads to strong cost of reproductive interference in the scarcest species limiting mimicry, mimicry is more likely to be observed between species of similar density (see Figure 4(f)).

Our results highlight that reproductive interference impacts the evolution of warning traits, and may even reverse the expected *model/mimic* relationship, depending on the relative abundances and individual defense levels of sympatric species.

## Discussion

### Reproductive interference alone cannot explain imperfect mimicry

Our results show that reproductive interference and historical constraints promoting ancestral traits can generate a continuous range of phenotypic distances from quasi perfect mimicry to warning trait displacement. Our study suggests that reproductive interference alone is unlikely to promote imperfect mimicry, in contradiction with previous predictions (Pfennig and Kikuchi, 2012). When sympatric species share the same ancestral trait, or in absence of historical constraints, we indeed observe either perfect mimicry or strong trait divergence, depending on the strength of reproductive interference. In our model, imperfect mimicry is observed only when stabilizing selection due to historical constraints promotes different ancestral traits in both species. The contrasted historical constraints undergone by the different species may thus play an important role in imperfect mimicry. These different constraints may be strongly correlated with the phylogenetic distances between species: closely-related species are likely to share similar genetic bases and developmental pathway leading to the warning trait and to also share similar environments, due to niche conservatism (Chazot et al., 2014; Elias et al., 2008; Joshi et al., 2017), likely to limit departure from ancestral trait values. Our results suggest that imperfect mimicry could not be promoted among closely-related species experiencing high levels of reproductive interference but limited differences in ancestral traits. Imperfect mimicry may rather be observed between phylogenetically-distant species, subject to more strikingly different historical constraints, where reproductive interference might be more limited. Distantly-related species indeed might have diverged in other traits, facilitating mate recognition through different cues.

For similar historical constraints, mimicry between defended species can then either be promoted or limited depending on predator discrimination. Low predator discrimination allows for the evolution of imperfect mimicry, since imperfect mimics are seen as similar by predators, allowing mutualistic relationship without implying strong cost of historical constraints. By contrast under high predator discrimination, mutualistic mimetic relationships necessitate a strong similarity between species, which is limited by historical constraints. Empirical studies based on vertebrates or on insects show that predators do not perceive difference between Batesian mimics and their models, or at least this difference does not entail a difference in behavior (Dittrigh et al., 1993; Kikuchi and Pfennig, 2010; Morris and Reader, 2016). Loose predator discrimination may therefore play a key role in the evolution of imperfect mimicry.

### How important are historical constraints in the evolution of warning trait?

Our model predict that imperfect mimicry can arise through the interplay between different historical constraints, predation, and reproductive interference. However, estimating the actual level of historical constraints potentially shaping the evolution of warning traits is not straightforward. Genetic studies, reviewed in Joron et al. (2006), show that *Heliconius* species share the same ‘toolkit’ of genes, repeatedly recruited during both convergent and divergent evolutions of warning traits within and between species. The important lability in color patterns observed in this genus suggests a limited level of developmental constraints, facilitating the evolution of mimicry, even between species from different sub-clades within this genus (Hines et al., 2011). Such mimicry between distantly related species shows that selection due to predation can overcome historical constraints. By contrast, in butterflies from the tribe Ithomiini and in butterflies from the tropical forests of the Western Ghats, India, a strong phylogenetic signal on the warning trait is observed (Chazot et al., 2014; Elias et al., 2008; Joshi et al., 2017), suggesting that historical constraints may limit the evolution of mimicry among distantly related species despite predation pressure.

### Evolution of female preferences limiting the reproductive interference costs generated by mimicry

When considering reproductive interference, the relationship between female and predator discriminations is crucial to understand the evolution of warning traits. Surprisingly, when female and predator discriminations are low, higher predator than female discrimination promotes convergence of warning traits because selection due to predation and reproductive interference increase with predator and female discrimination respectively. By contrast, when female and predator discriminations are high, imperfect mimicry can evolve despite reproductive interference. When female discrimination is high, successful species recognition might occur without decreasing the protection brought by mimicry. Such situation arises when predators largely generalize, and therefore do not discriminate imperfect mimics. Some studies report similar female and predator discriminations (Finkbeiner et al., 2014; McClure et al., 2019), suggesting that reproductive interference may act on mimetic species. On the other hand, differences in the discrimination of color patterns between prey and predators may exist in the wild. For instance, Llaurens et al. (2014) showed that the variations in color pattern between co-mimetic species from the distantly related genera *Heliconius* and *Melinaea* might be better perceived by the *Heliconius* butterflies themselves but not by avian predators. The evolution of visual perception in females could also enhance species discrimination without impairing mimicry. The evolution of vision in females from the *Heliconius* butterflies indeed coincides with the evolution of the yellow pigments 3-OH-kinurenin displayed on their wings (Bybee et al., 2012). The evolution of high discrimination capacities in mimetic prey, as well as the evolution of mating cues undetected by predators could thus limit the cost of reproductive interference in mimetic prey. In butterflies, mate choice indeed often relies on pheromones that may strongly differ among closely-related mimetic species (Darragh et al., 2017; González-Rojas et al., 2020). Similarly, in non-mimetic species, chemical cues may reduce reproductive interference without entailing reproductive character displacement on a trait under natural selection. Females of the swordtails *Xiphophorus pygmaeus* prefer larger mate leading to reproductive interference with males of the *Xiphophorus nigrensis* species. However *X. pygmaeus* females avoid mating with heterospecific on the basis of chemical cues (Crapon de Caprona and Ryan, 1990). Micro-habitat differences among mimetic species may also allow reducing heterospecific encounters, while still benefiting mimicry by sharing the same predator community (Estrada and Jiggins, 2002). For example the two sympatric ladybird species *Harmonia axyridis* and *Harmonia yedoensis* have similar body size and coloration (Sasaji, 1998) and experience reproductive interference (Noriyuki et al., 2012). These species nevertheless have different host specialization (Noriyuki et al., 2011), that may limit reproductive interference (Noriyuki, 2015). Likewise, in three *Morpho* butterfly species displaying local convergence in wing patterns (Llaurens et al., 2021), temporal segregation in patrolling activity has been observed between species sharing similar color patterns (Le Roy et al., 2020), which may strongly limit heterospecific rivalry.

The levels of reproductive interference among mimetic species might thus be modulated by the evolution of the converging traits themselves, as well as the evolution of other traits involved in species interactions.

### Reproductive interference strongly impacts species with low relative density

Our model shows that the effect of reproductive interference strongly depends on the relative abundances of interacting species, leading to surprising evolutionary outcomes. For example, in rare defended species, selection favoring mimicry towards a defended *model* species is expected to be strong. Nevertheless, our model shows that an elevated cost of reproductive interference prevents the evolution of mimicry in the rarest species, because females then encounter much more heterospecific than conspecific males.

Reproductive interference may particularly promote the emergence and persistence of a distinct warning trait in low-density populations of warning species coming into contact with a local mimicry ring that exhibits a different warning trait. Our model does not take into account the dynamics of population density, and therefore ignores the extinction risk of low-density populations. Such non-mimetic populations with low density might nevertheless persist in the wild, when the level of individual defense is sufficiently high.

Because undefended mimics have a negative impact on predator learning (Lindström et al., 1997), they are expected to be scarce compare to their models (Kunte et al., 2021). In line with this prediction, empirical studies report low density of undefended mimics compared to their defended models (Long et al., 2015; Prusa and Hill, 2021). Reproductive interference may act strongly on Batesian mimics because of their low density with respect to the *model* species. Our model thus suggests that Batesian mimicry among closely related species may be limited by strong reproductive interference acting on Batesian mimics due to phylogenetic proximity and unbalanced density. By contrast, Batesian mimicry may evolve between distantly-related species despite unbalanced density, because high phylogenetic distance reduces risk of reproductive interference. This is supported by the pattern of convergence observed in tropical forests of the Western Ghats in India: Müllerian mimicry is observed between closely related species whereas Batesian mimicry involves more distantly related species (Joshi et al., 2017).

Reproductive interference does not always promote divergence of reproductive character (here the warning trait) but can also provoke spatial segregation between species (Gröning and Hochkirch, 2008). The strong reproductive interference acting on scarce mimetic species may limit their coexistence with more abundant mimetic species displaying similar warning signals. Reproductive interference may then restrict the spatial distribution of mimetic species with low abundance to the edges of the range occupied by more abundant co-mimetic species.

Our model also brings new insights to the ecological processes driving the direction of advergence in warning traits. In the absence of reproductive interference, mimicry is expected to evolve in the less defended species (*e.g*. low density populations and/or low level of individual defense) towards better-defended species living in sympatry (Balogh and Leimar, 2005; Franks and Sherratt, 2007). When considering the cost of reproductive interference, however, this general trend does not always hold. Our results show that warning traits in the most defended species can evolve toward the warning trait of the less abundant one.

Our model therefore highlights the interplay between mutualism and reproductive interference in sympatric species, which determines the strength and the direction of traits evolution involved in these ecological interactions.

### Reproductive interference can explain the emergence of mimetic diversity

In our model, we consider the evolution of warning trait between two interacting species. In the wild however, natural communities involve a variable number of mimetic species, with mimicry ring size ranging from 2 to a dozens of species (Kunte et al., 2021). Assuming reproductive interference, species richness within a mimicry ring may influence warning trait evolution. We hypothesize that mimicry rings with high species richness are more likely to contain closely related species. The evolution of mimicry in a species may be limited if a closely related species is abundant in the ring because of the strong cost of reproductive interference generated. However, in a mimicry ring with high species richness, distantly-related species may not suffer from less reproductive interference but mutually benefit of sharing a similar warning trait. This increased advantage of mimicry between distantly-related species may counterbalance the more elevated costs due to reproductive interference with closely-related species, increasing the likelihood of having closely related species within large mimicry rings.

Our results shed light not only on the persistence of distinct warning traits within local communities of defended species in the wild, but also on the emergence of these distinct warning traits in the first place. Mimetic diversity is an apparent paradox but several hypotheses have been suggested to promote the persistence of different warning signals, such as the segregation of predators within microhabitats (Beccaloni, 2008; Devries et al., 1999; Elias et al., 2008; Willmott et al., 2017). The spread of distinct warning traits has frequently been shown to be promoted by demographic stochasticity, as in shifting balance models (Mallet and Joron, 1999; Sherratt, 2006) or in other models combining predator behaviors, such as neophobia, to stochastic effects (Aubier and Sherratt, 2015). However, these models do not provide any selective mechanism explaining the emergence of warning signal with different levels of divergence, contributing to mimetic diversity. By contrast, reproductive interference selects for different levels of divergence in warning traits, and could be a major driver of the diversity of mimetic traits. Other mechanisms may generate gradual departure from the ancestral trait value and may also contribute to the diversity of mimetic traits: the evolution of aposematic signals in defended species away from those exhibited in Batesian mimics has been theoretically shown (Franks et al., 2009), but empirical evidence of such effect of Batesian mimicry is still lacking. Artificial modification of the warning trait of mated females has also been demonstrated to reduce harassment by males in the butterfly *H. erato*, and would therefore allow them to lay more eggs, suggesting that evolution of slightly divergent trait could be promoted in females (Merrill et al., 2018).

We hope our theoretical work will encourage experimental approaches investigating the impact of reproductive interference on mimicry. Such studies may shed lights on the actual role of reproductive interference on mimetic diversity.

## Conclusion

Our analytical and numerical results show that reproductive interference and historical constraints can explain a wide range of levels of convergence, and even explain divergence of warning trait between sympatric species. Our results suggest that reproductive interference alone cannot explain imperfect mimicry, highlighting the role of historical constraints in the evolution of imperfect mimicry. Our study also highlights the importance of female and predator discriminations in the evolution of warning traits.

## Data Availability

Codes are available online: github.com/Ludovic-Maisonneuve/limits-of-ev-conv

## Acknowledgments

LM would like to thank Dorian Ni for feedbacks on the mathematical part of the study. LM and VL would like to thank Charline Pinna and the whole ‘Evolution and Development of Phenotypic Variations’ team for stimulating discussions on the evolution of warning traits. The authors would like to thank the ANR SUPERGENE (ANR-18-CE02-0019) for funding the PhD of LM, and the Emergence program from Paris city council for supporting the team of VL. This work was also supported by the Chair “Modélisation Mathématique et Biodiversité” of VEOLIA-Ecole Polytechnique-MNHN-F.X.

## Appendix: The limits of evolutionary convergence in sympatry: reproductive interference and historical constraints leading to local diversity in aposematic signals

In section 1 we detail the hypothesis allowing to approximate the evolutionary dynamics. In Section 2 we detail how some components of fitness can be approximated using that the genetic covariances of traits and preference are low. In Section 3 we detail the computation of the selection vector describing the impact of the selective forces on the evolution of traits and preference. In Section 4, in line with the weak selection hypothesis we suppose that the genetic correlation between trait and preference is at equilibrium and estimate this quantity.

In Section 5 we first consider the classical case where the species 1 is a *mimic* of the *model* species 2. We thus assume higher abundance (*n*_2_ ≫ *n*_1_) and greater defense level in species 2 as compared to species 1. In this case, species 1 weakly impacts the evolution of trait and preferences in species 2, neither through mimicry nor through reproductive interference. We then assume fixed mean trait and preference in species 2, sticking to the ancestral trait value 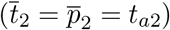.

In Section 6 we then study the joint evolution of trait and preference in both species 1 and 2, accounting for the general case of mimicry between sympatric species with various relative abundances and defense levels, where evolution of traits can occur in both species.

### 1 Approximation of the evolutionary dynamics

When the variance of trait and preference is small relative to the curvature of the fitness function in each species we get the approximation of the evolutionary dynamics in Equation (7).

We assume weak natural and sexual selective pressures: *a*_1_, *a*_2_, *d, b* and *c_RI_* are of order *ε* and *s*_1_ and *s*_2_ are of order *ε*^2^, where *ε* is little. Assuming selective coefficients *s*_1_ and *s*_2_ to be of order *ε*^2^ allows to explore the interplay between historical constraints, predation and reproductive interference when selective pressures due to different evolutionary forces are of the same order of magnitude (see Appendix 2). Note that depending on parameters value we can also explore cases where an evolutionary force dominate the others.

Under weak natural and sexual selective pressures the approximation in Equation (7) stands. Moreover it allows to estimate the genetic covariance between trait and preference in both species. We can then analytically determine the mean traits and preferences values at equilibrium in most cases. In the few remaining cases, we use numerical analyses to estimate these values.

Because the stringency of the discrimination behavior of female and predator are key drivers of the advergence toward the trait displayed in species 2, we then explore larger values of female and predator discriminations, *i.e. a*_1_ = *O*(1) and *b* = *O*(1). Under such assumptions, the effects of predation and reproductive interference on the changes in the mean trait and preference are of order *ε*. We also assume that *s*_1_ is of order *ε*, because if *s*_1_ were of order *ε*^2^ the strength of historical constraints would always be negligible as compared to the other two selective forces. Assuming strong female discrimination violates the weak selection hypothesis, because strong female discrimination generates strong sexual selection for males and large opportunity cost for females. However, because strong female discrimination leads to higher sexual selection, the discrepancy between preference and trait values 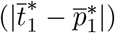 becomes limited. Therefore, sexual selection and opportunity cost are actually weak and we can still estimate the matrix of genetic covariance and assume that the genetic variances of traits and preference are low. We assume that the variances of trait and preference are low to keep them small relative to the curvature of the fitness function in each species when (*a*_1_ = *O*(1) and *b* = *O*(1)). However, because of the complexity of the model, under strong female and predator discriminations (*a*_1_ = *O*(1) and *b* = *O*(1)), we cannot analytically determine the mean trait and preference values at equilibrium (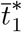 and 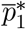). We then use numerical simulations of Equation (**??**) to explore cases where *a*_1_ and *b* are of order 1 and *s*_1_ is of order *ε*.

### 2 Relative small variance approximation

Because we assume that the variance of traits and preferences is small relative to the curvature of the fitness function we can approximate Equations (5) and (6). Here we detail how we obtained these approximations. 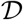 is defined by

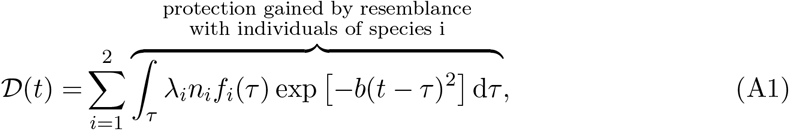

Let *i* ∈ {1, 2}, we have

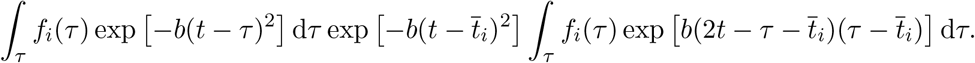

Using a Taylor expansion of exp 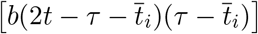 we have

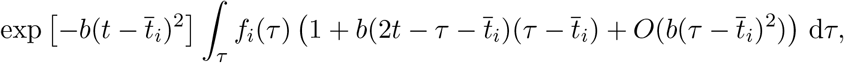

which is equal to

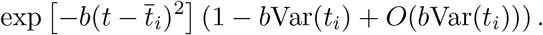

Hence when the variance of *t_i_* is small relative to the curvature of the fitness function (implying than *b*Var(*t_i_*) is low) 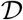 can be approximated by

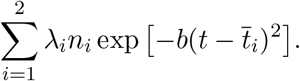

The reasoning is similar to approximate 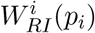 so for *i* ∈ {1, 2}:

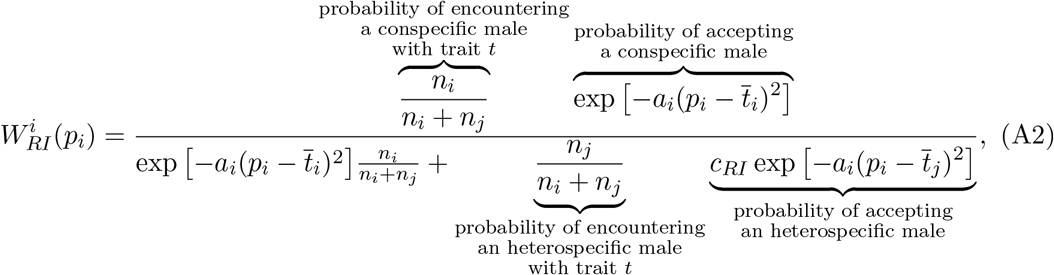

with *j* ∈ {1, 2} with *j* ≠ *i*.

### 3 Selection coefficients

In this section we detail how we obtain the expressions for the selection vectors.

For *i* ∈ {1, 2}, the selection coefficients are defined by:

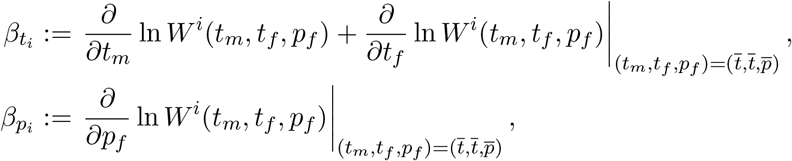

where *W^i^* are the fitness of a mated pair of a male with trait *t_m_* and a female with trait *t_f_* and preference *p_f_* of species *i*, and write

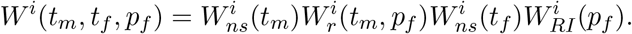

Here 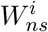 is the impact of natural selection on fitness and 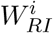 is the impact of reproductive interference on fitness of individuals of species *i*.

#### 3.1 Computation of *β*_*t*_1__ and *β*_*t*_2__

For *i* ∈ {1, 2} the selective forces acting on the trait value in species *i* write:

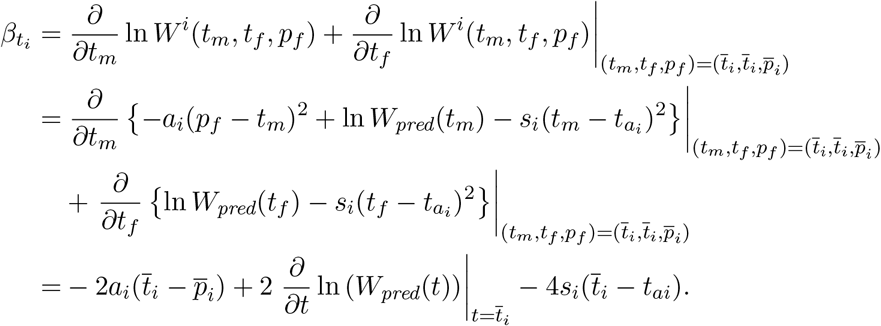

Recall the definition of *W_pred_* in (10). The second term of the right hand side equals

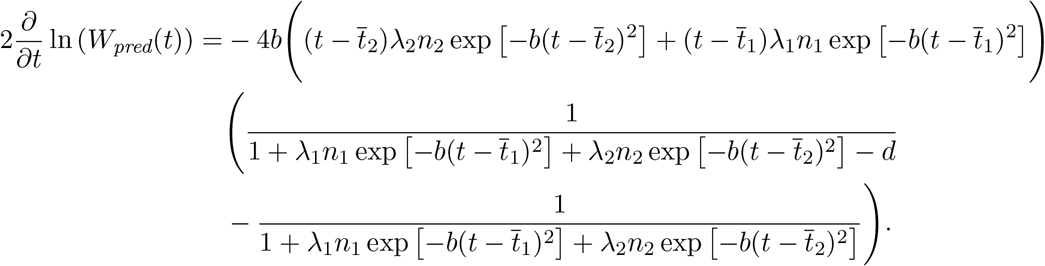

When the parameters *b* and *d* are of order *ε*, we obtain

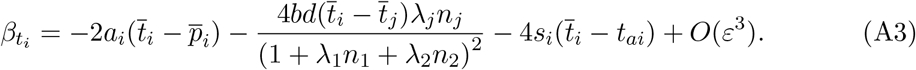

#### 3.2 Computation of *β_pi_*

For *i* ∈ {1, 2} the selective forces acting on the preference value in species *i* write:

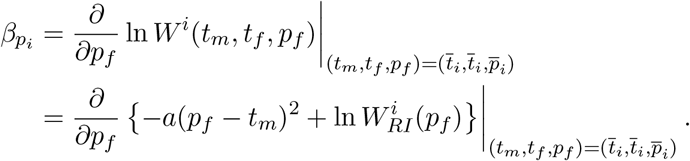

The last term of the right hand side equals

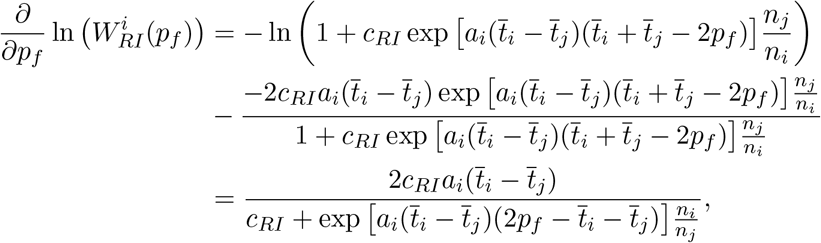

with *j* ∈ {1, 2}, *j* ≠ *i*.

When the parameters *a*_1_, *a*_2_ and *c_RI_* are of order *ε*, we finally obtain

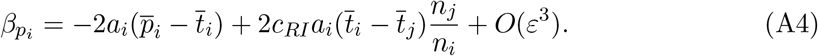

### 4 Computation of the matrix of correlation

In this part we approximate the genetic covariance between trait and preference *C_t_i_p_i__* in species *i* for *i* ∈ {1, 2}, using the results from [Kirkpatrick et al., 2002]. Trait and preference are controlled by different sets of unlinked loci with additive effects, denoted *T_i_* and *P_i_*, respectively. For each *j* in *T_i_* (resp. *P_i_*), we note 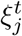 (resp. 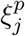) the contribution of the locus *j* on trait (resp. preference) value. The trait and preference values of an individual are then given by

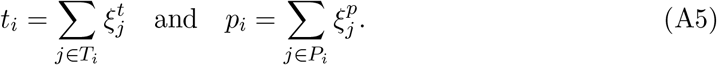

As in [Lande, 1981] we assume that the distributions of 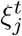 and 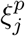 are multivariate Gaussian. Let *C_jk_* be the genetic covariance between loci *j* and *k*. Then the elements of the matrix of correlation are given by:

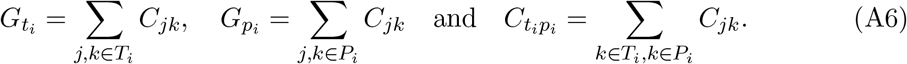

To compute the change on genetic correlation we need to identify various selection coefficients (see [Barton and Turelli, 1991, Kirkpatrick et al., 2002]). These coefficients are obtained using the fitness of a mated pair of a male with trait *t_m_*, and a female with trait *t_f_*, and preference *p_f_ W^i^*(*t_m_, t_f_, p_f_*), defined in Equation (1).

For simplicity we consider only leading terms in the change in genetic correlation. For (*j, k*) ∈ *T_i_* × *P_i_*, combining Equations (9), (12), (15) from Kirkpatrick et al. [2002] gives the change in the genetic covariance between loci *j* and *k*:

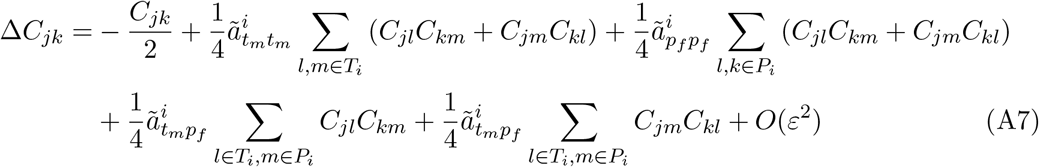

with 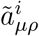 for (*μ, ρ*) ∈ {*t_m_, t_f_, p_f_*}^2^ being the leading term of

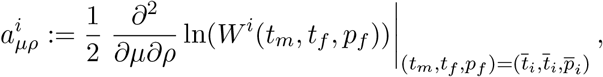

the selection coefficient calculated from the fitness of a mated pair of a male with trait *t_m_* and a female with trait *t_f_* and preference *p_f_*. The expressions of these coefficients are:

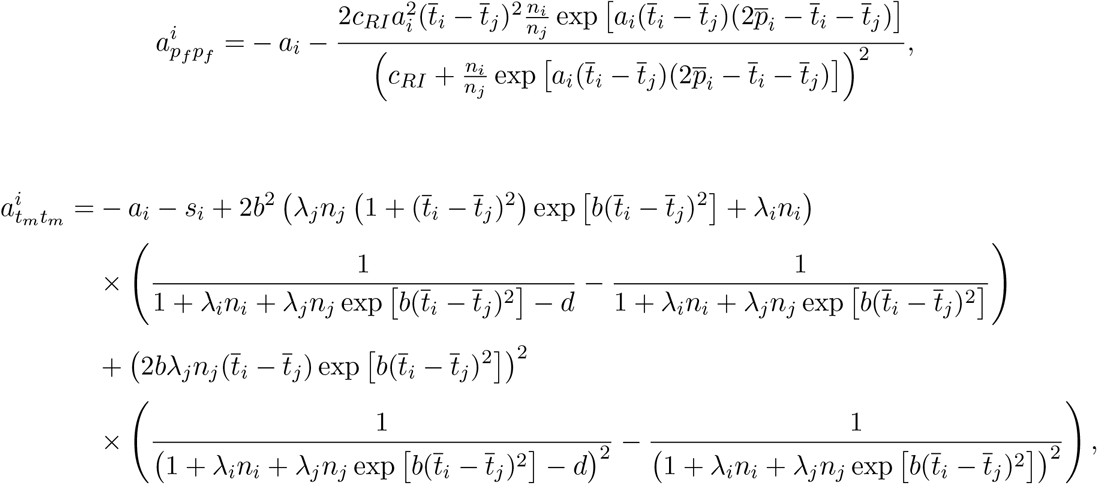

and

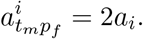

A Taylor expansion gives 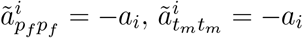 and 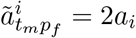.

By summing Equations (A7) over each *j, k* in *T_i_* and *P_i_* we obtain:

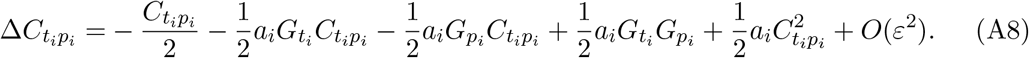

For the sake of simplicity we assumed that the genetic correlations between traits and preference are at equilibrium (as in [Barton and Turelli, 1991, Pomiankowski and Iwasa, 1993]). We obtain from (A8) that the value at equilibrium is given by

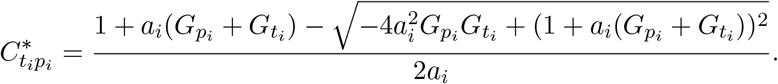

Because the genetic variances of trait and preference are low we have

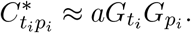

When we relax the hypothesis *a_i_* = *O*(*ε*) and *b* = *O*(*ε*) the previous calculation still stands, except that 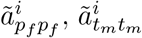 and 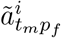 are of order 1 instead of being of order *ε*.

### 5 Evolution of warning trait toward a fixed defended species

We assume higher abundance in species 2 compared to species 1 (*n*_2_ ≫ *n*_1_). We also assume a great defense level in species 2. In this case species 1 weakly impacts the evolution of trait and preferences in species 2 through mimicry and reproductive interference. We assume fixed mean trait and preference in species 2 equal to the ancestral trait value 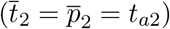. This hypothesis, which we will relax later, allows a full analytical resolution (see below).

In Section 4.1 we compute the values of trait and preference that cancel the leading term of the selection vector. In Section 4.2 we study when trait and preference converge toward the point found in the previous section. In Section 4.3 we study the impact of weak or strong female discrimination hypothesis on the phenotypic distance between the two species. In Section 4.4 we detail the impact of each parameter of the phenotypic distance between the two species.

#### 5.1 Quasi equilibria

We search mean values of trait and preference at equilibrium 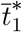 and 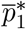 such as the leading terms of 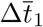 and 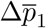 are null. We remind that we assume weak female and predator discriminations. The leading terms of 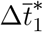 and 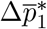 are thus 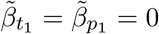 with

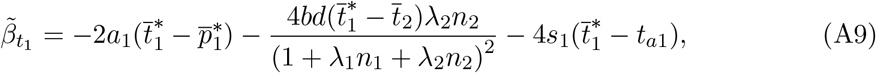

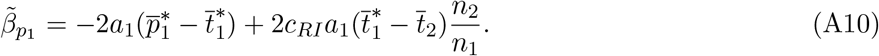

We call such points quasi equilibria because they are not equilibrium point of Equation (11) but only cancel the term of leading order.

##### Lemma 1.

*If*

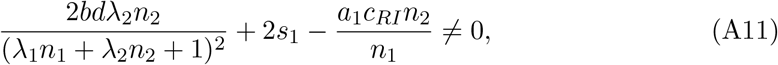

*there is one quasi equilibrium point* 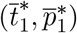, *where*

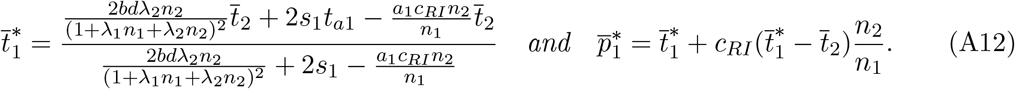

*Proof*. Assume that *a*_1_ ≠ 0. From the definitions of 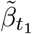 and 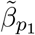 in (A9) and (A10), we deduce that the values of trait and preference 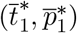 for which the selections on trait and preference are null satisfy:

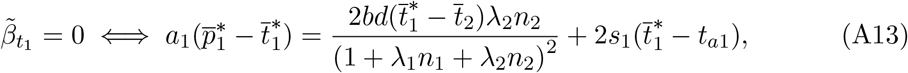

and

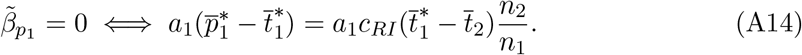

From (A13) and (A14) we derivate:

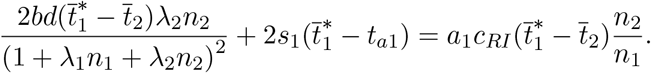

By factorizing by 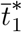 we get:

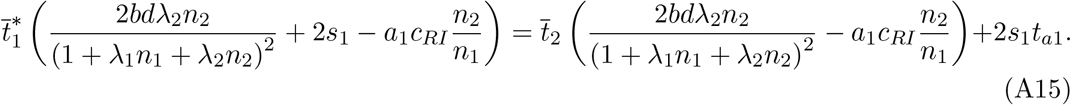

Using (A11), we obtain the value of 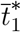 given in (A12). From (A14) we deduce the associated mean preference and conclude the proof.

#### 5.2 Fast and slow dynamics

In this section we study when trait and preference in species 1 converge towards the quasi equilibrium point.

We suppose that the parameters are such as 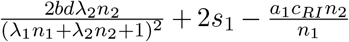 is not of an order inferior to *ε*^2^. We may have two long term behaviours. In case of convergence (case 1. of Lemma 2), the trajectories quickly approach the line 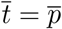, before evolving according to a slower dynamic along this line (see the proof of Lemma 2 and Figures A1 and A2).

##### Lemma 2.

*1. If*

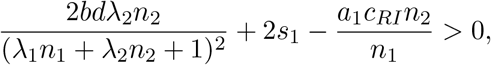

*the quantities* 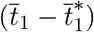 *and* 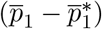 *become of order 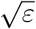 when the number of generations goes to infinity, where the values of 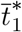 and 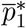 have been given in* (A12).

*2. If*

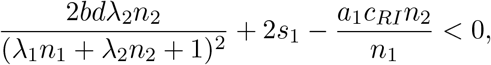

*trait and preference become very large*.

*Proof*. As we have said, we can then decompose the dynamics into two steps. In the first one the leading order terms of the selection coefficient are the terms describing sexual selection and cost of choosiness. In this case we can approximate 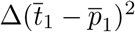.

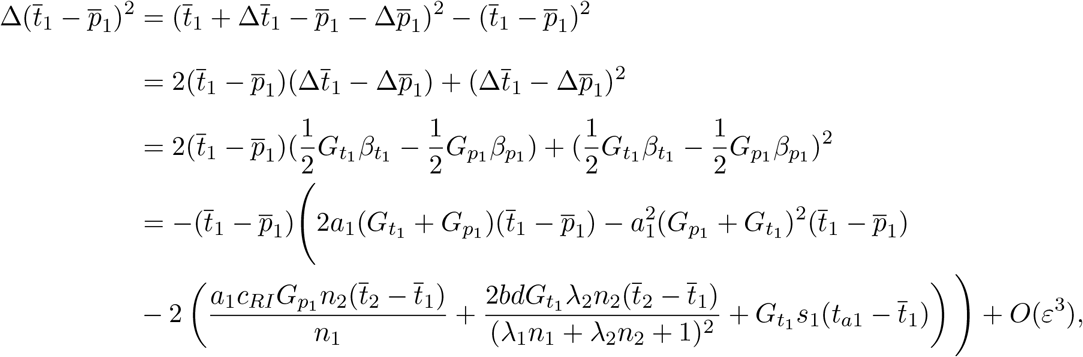

where we used (11) for the third equality and (A3) and (A4) for the last one. As by assumption, *a*_1_, *c_RI_, b* and *d* are of order *ε*, and *s* of order *ε*^2^, we obtain

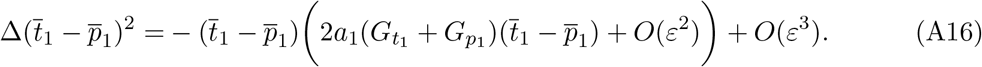

Denote by *r_n_* the value of 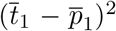 at generation *n*. We will prove the following statement:

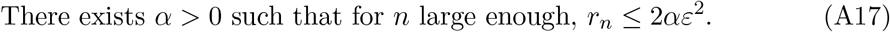

If we introduce the parameter

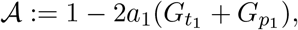

Equation (A16) may be rewritten:

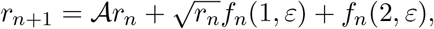

for every 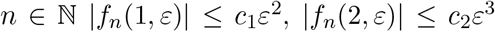 where *c*_1_ and *c*_2_ are finite constants.

Now introduce the variable

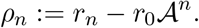

As *a*_1_ is of order *ε*, and *G*_*t*_1__ and *G*_*p*_1__ of order one, there exists a positive and finite constant 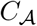 such that for *ε* small enough,

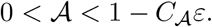

We will choose in the sequel a positive real number *α* satisfying

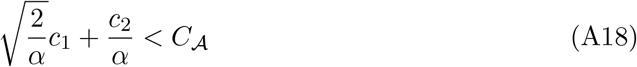

and

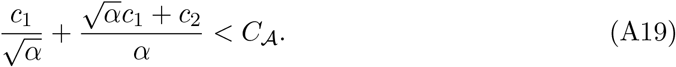

As 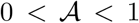, we see that for *n* large enough, 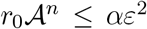. Thus to prove (A17), it is enough to prove that for *n* large enough, |*ρ_n_*| ≤ *αε*^2^. We obtain for *ρ* the recurrence equation

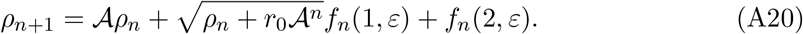

Assume first that |*ρ_n_*| ≤ *αε*^2^. In this case,

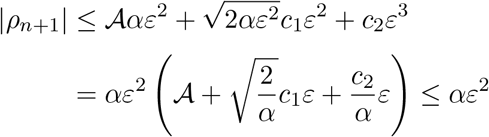

thanks to (A18) provided that *ε* is small enough. Hence it is enough to prove that there exists one *n* such that *p_n_* ≤ *αε*^2^, and the inequality will hold for later generations.

Now assume that *p_n_* > *αε*^2^. Using that

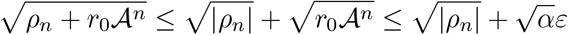

for *n* large enough, we obtain from (A20)

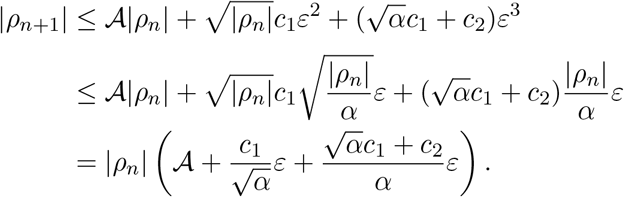

Due to (A19), the term in bracket belongs to (0, 1) if *ε* is small enough. We conclude that there exists *n*_0_ such that |*p_n_*| ≤ *αε*^2^ for any *n* ≥ *n*_0_. This concludes the proof of (A17) and of the fast convergent phase.

We now want to prove that 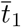 becomes close to 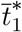, and 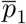 to 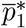. To this aim, we look at the variation of

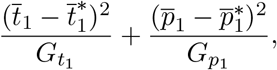

where the values of 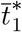 and 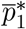 are the ones defined in (A12). By definition,

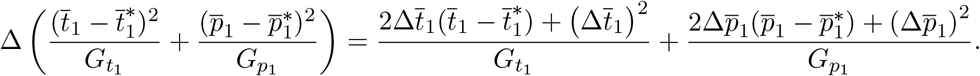

When 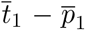 is of order *ε*, 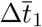 and 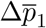 are of order *ε*^2^. We then neglect the terms proportional to 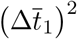 and 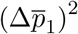. This yields

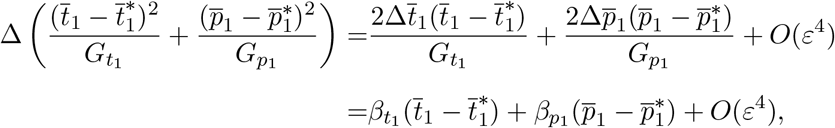

where we used (11). Now notice that

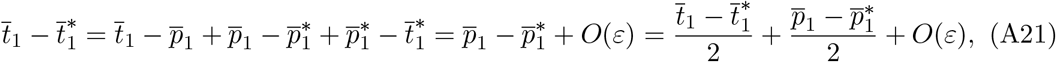

as 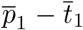 is of order *ε* and 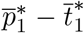 of order *ε*. We can make the same computation for 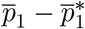. Adding that *β*_*t*_1__ and *β*_*p*_1__ are of order *ε*^2^, we obtain:

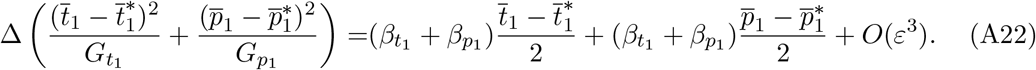

Using (A3) and (A4) and simplifying, we get:

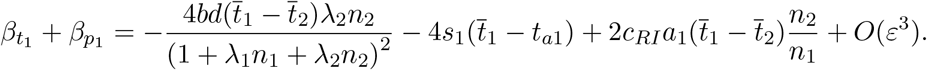

By substracting 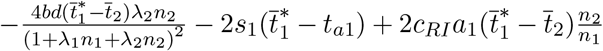 which is equal to zero (because 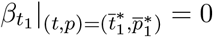 and 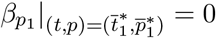 we get:

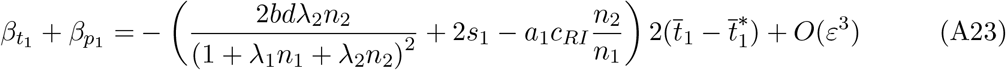

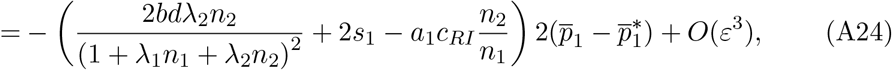

where we used (A21). Replacing *β*_*t*_1__ + *β*_*p*_1__ using (A23) and (A24) into (A22) yield:

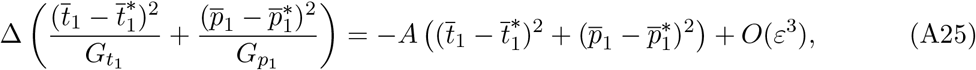

with

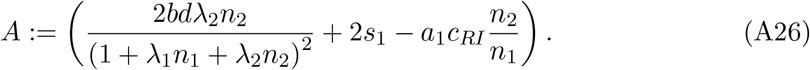

**First case:** *A* > 0

We suppose that the parameters are such that A is not of an order inferior to *ε*^2^. Using computations similar to the ones derived to prove (A17), we may prove that after a number of generations large enough, 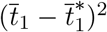 and 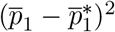 become of order *ε*, and thus 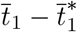 and 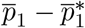 become of order *ε*^1/2^.

To illustrate this result we simulated two trajectories of the system (11) 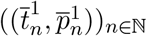 and 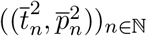 for two different initial conditions: (*t*^*i*1^, *p*^*i*1^) and (*t*^*i*2^, *p*^*i*2^). The trajectories are shown in Figure A1 and the parameter values are given in the caption. The parameter values correspond to the weak selection hypothesis and are such that *A* > 0.

The fast and slow dynamics are visible in Figure 1(a) where the trajectories quickly become close to the line defined by 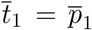 and then slowly converge toward the quasi equilibrium points. We noted 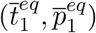 the equilibrium point reached by the two simulated trajectories.

**Figure A1:**
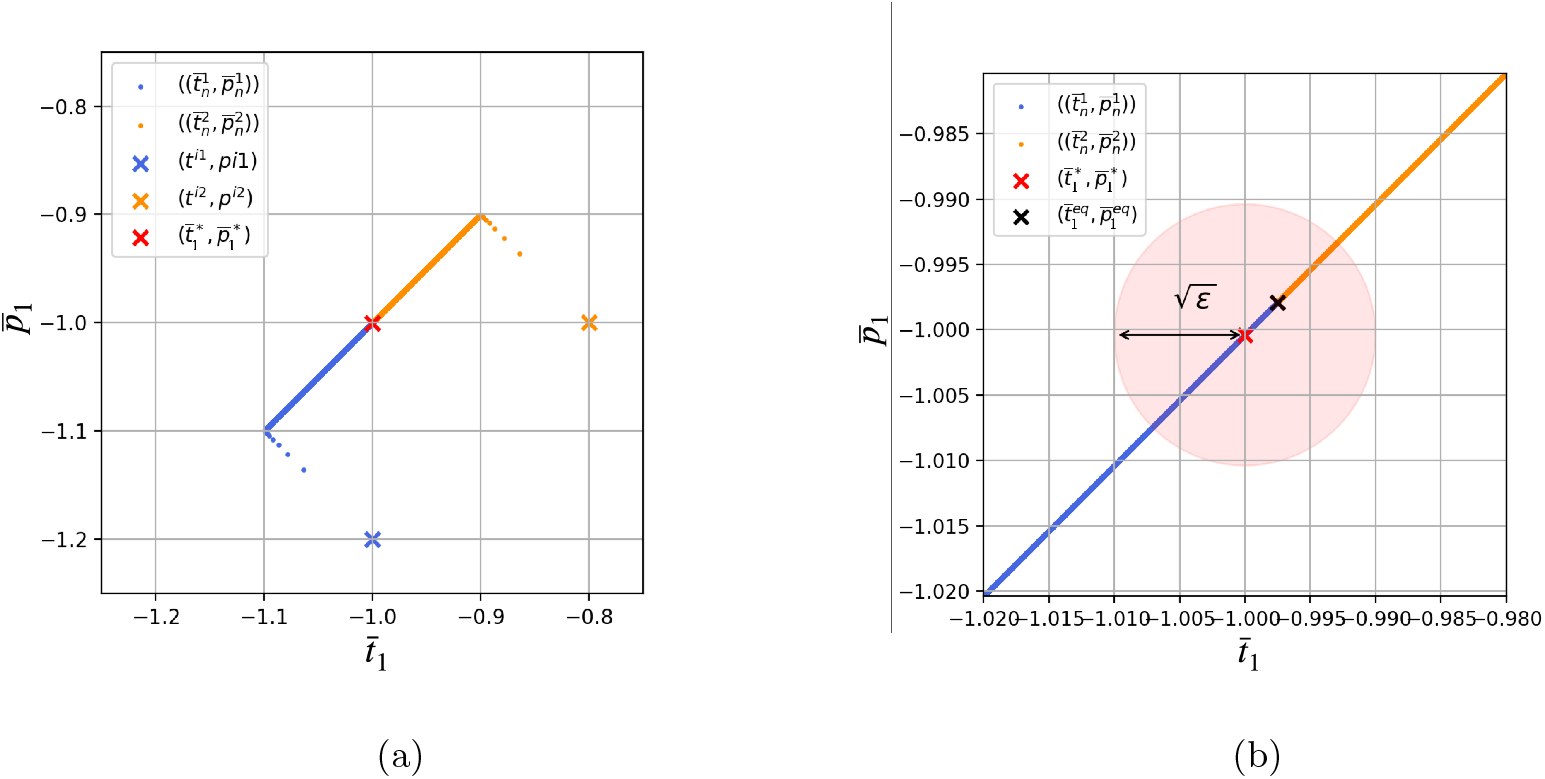
(a) Evolution of mean trait value 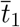 and mean females preference value 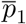 for two different initial conditions. The blue (resp. orange) points represent the numerical trajectories for the initial condition (*t*^*i*1^, *p*^*i*1^) (resp. (*t*^*i*2^, *p*^*i*2^)) (b) Zoom on the previous figure. The red circle has for center the quasi equilibrium point 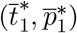 and for radius 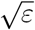. This illustrates that the numerical steady point 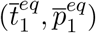 is at a distance of order 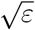. The trajectories were obtained with the parameter values: (*t*^*i*1^, *p*^*i*1^) = (−1, −1.2), (*t*^*i*2^, *p*^*i*2^) = (−0.8, −1), *G*_*t*_1__ = *G*_*p*_1__ = 0.01, *a*_1_ = 10^-4^, *b* = 2 × 10^-4^, *d* = 2 × 10^-4^, *λ*_1_ = 0.1, *λ*_2_ = 0.1, *n*_1_ = 10, *n*_2_ = 20, *c_RI_* = 10^-4^, *s*_1_ = 10^-8^, *t*_*a*1_ = 0 and *t*_*a*2_ = 1. The trajectories are computed over 1,000,000 generations.

**Second case:** *A* < 0.

Recall (A25) and that we assumed that A is not of order inferior to *ε*^2^. This entails that 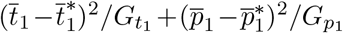 goes to infinity. However when 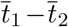 or 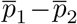 becomes high (for example of order 1/*ε*) the approximation made in section 2 does not stand. Therefore from a certain number of generations the approximation does not correctly describe the evolution of trait and preference. We can only conclude that trait and preference become very large. This is illustrated by Figure A2 which shows a simulation of two trajectories when *A* < 0.

**Figure A2:**
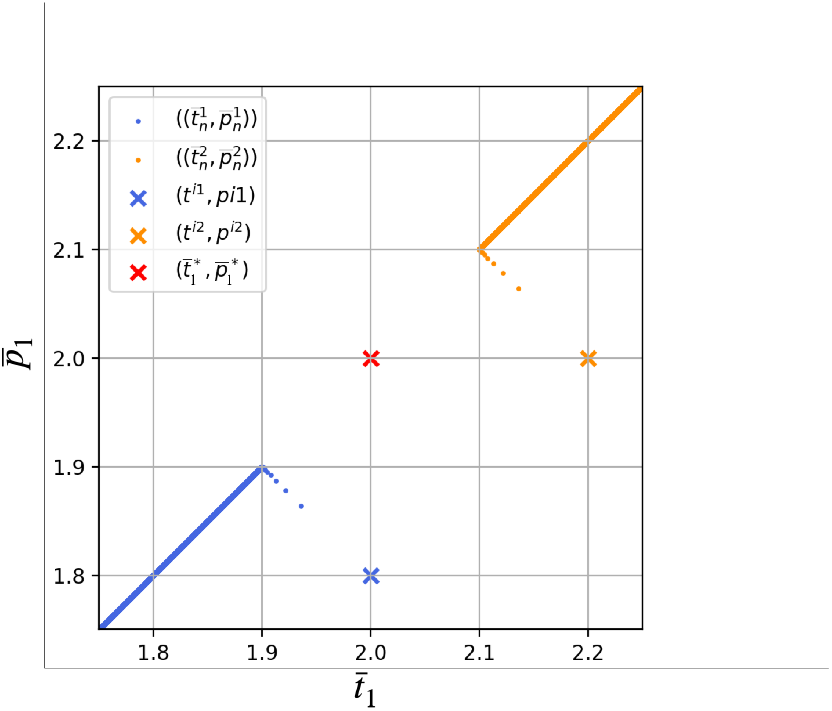
Evolution of mean trait value 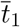 and mean females preference value 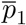 for two different initial conditions. The blue (resp. orange) points represent the numerical trajectories for the initial condition (*t*^*i*1^, *p*^*i*1^) (resp. (*t*^*i*2^, *p*^*i*2^)). The trajectories were obtained with the parameter values: (*t*^*i*1^, *p*^*i*1^) = (−1, −1.2), (*t*^*i*2^, *p*^*i*2^) = (−0.8, −1), *G*_*t*_1__ = *G*_*p*_1__ = 0.01, *a*_1_ = 10^-4^, *b* = 2 × 10^-4^, *d* = 2 × 10^-4^, *λ*_1_ = 0.1, *λ*_2_ = 0.1, *n*_1_ = 10, *n*_2_ = 20, *c_RI_* = 10^-4^, *s*_1_ = 0.25 × 10^-8^, *t*_*a*1_ = 0 and *t*_*a*2_ = 1. The trajectories are computed over 1,000,000 generations.

#### 5.3 Effect of weak or strong female discrimination on the phenotypic distances between the two species

Previously we obtained an analytic approximation of the phenotypic distance between the two species when the strengths of female and predator discriminations (*a*_1_ and *b*) were weak (i.e. *a*_1_ = *O*(*ε*) and *b* = *O*(*ε*)). To obtain this approximation we also assumed that the strength of historical constraints has the same order of magnitude as selection coefficients due to predation and reproductive interference (i.e. *s*_1_ = *O*(*ε*^2^)). Under this assumption, the phenotypic distance between the two species becomes very large (see Figure A3) when

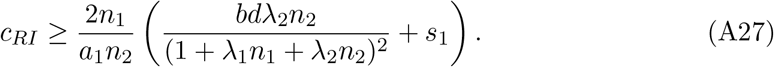

As explained in the main manuscript, this unrealistic divergence (under the weak female discrimination hypothesis (*a*_1_ = *O*(*ε*))) suggests that reproductive interference promotes strong phenotype divergence between the two species. Under weak female discrimination, only a large distance between females preference and the trait displayed by species 2 leads to a reduction of fitness cost generated by reproductive interference. Because females preference generates sexual selection on males trait, the phenotype divergence between the two species becomes high. When we relax the weak female discrimination hypothesis and use numerical simulations, then a great cost of reproductive interference provokes large but still of order 1 phenotypic distance between the two species (see Figure A3). However, we observe that the lower female discrimination, the higher is phenotypic distance when the cost of reproductive interference increases. As explained above, when females are not very choosy, reproductive interference favors the evolution of larger phenotypic distances limiting heterospecific mating.

**Figure A3:**
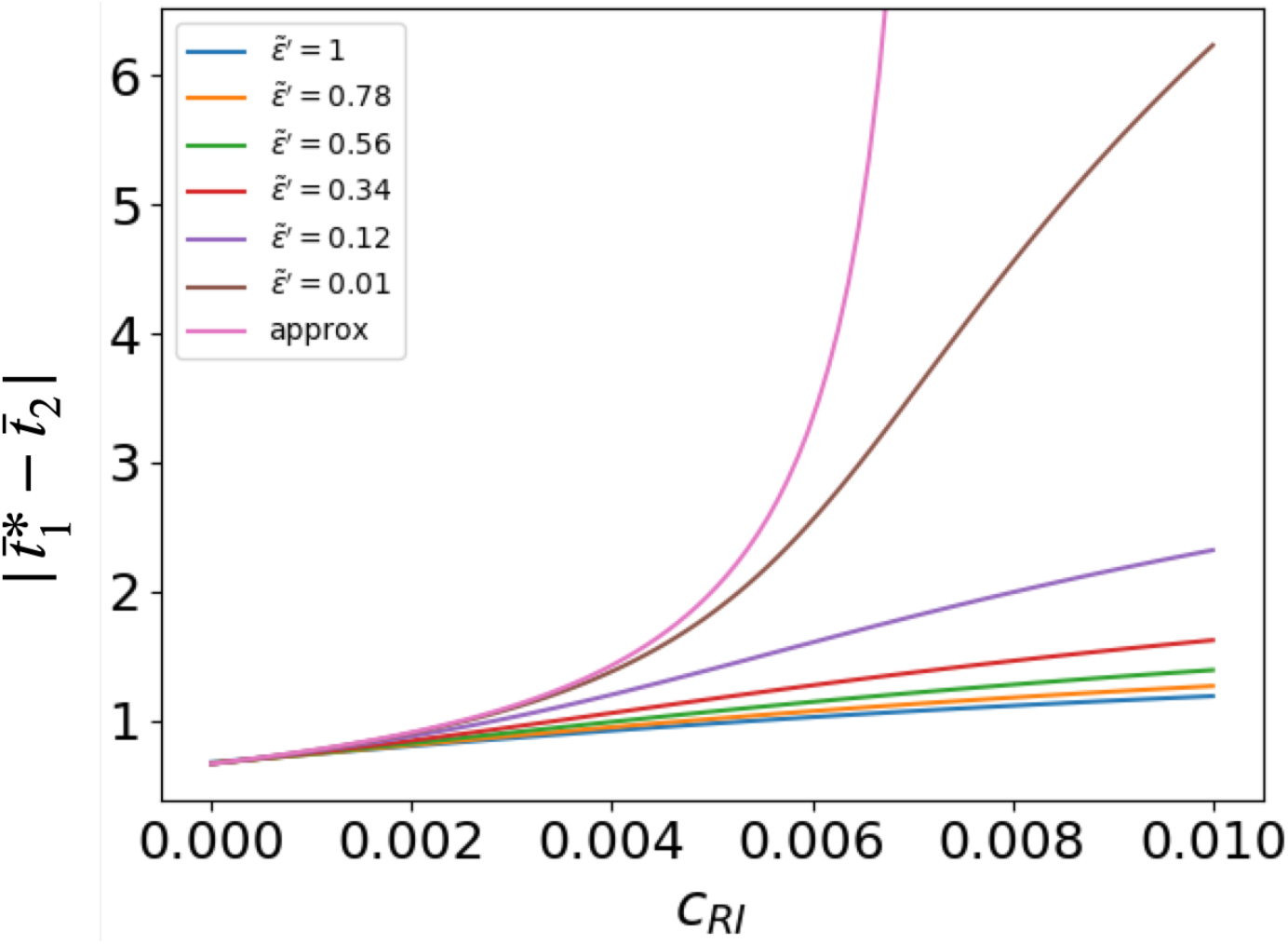
Influence of the cost generated by reproductive interference *c_RI_* on the phenotypic distances between the two species, using the analytical approximation (‘approx’ curve) or numerical simulations. Different values of female and predator discriminations coefficient (*a*_1_ and *b*) and strength of historical constraints (*s*) are assumed for numerical simulations. This illustrate that curves obtained by numerical simulations tend to look similar to those using the analytical approximation where parameters values tends to satisfy weak female and predator discriminations. We assume: *G*_*t*_1__ = *G*_*p*_1__ = 0.01, 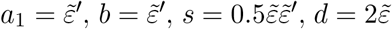, *λ*_1_ = 0.1, *λ*_2_ = 0.1, *n*_1_ = 10, *n*_2_ = 20, *t*_*a*1_ = 0 and *t*_*a*2_ = 1.

#### 5.4 Effect of different parameters on the level of trait divergence 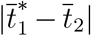

We then aim at disentangling the effects of the different parameters of the model on trait divergence. We thus focus on the case where *A* > 0 (recall definition (A26)). In this case, assuming weak female and predator discriminations, we know that the phenotypic distance between the two species, 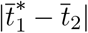, equals

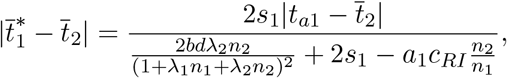

allowing to investigate the relative impact of the different parameters on trait divergence. We also run numerical simulations to investigate the effect of the different parameters assuming strong females preference and predator discrimination (*i.e. a*_1_ = *O*(1) and *b* = *O*(1)).

##### 5.4.1 Effect of reproductive interference

Assuming weak female and predator discriminations, the impact of the strength of reproductive interference on the phenotypic distance between the two species is given by the sign of 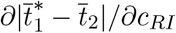. In our case,

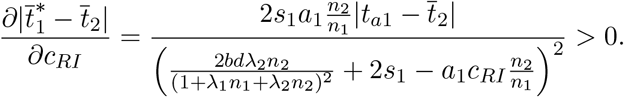

Hence the level of divergence increases with the strength of reproductive interference. A similar effect is observed assuming strong female and predator discriminations (see Figure A4(a)). Similarly, assuming weak female and predator discriminations

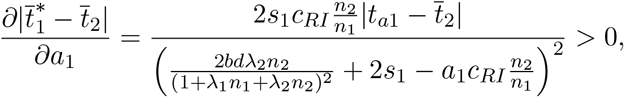

thus the phenotypic distance (along with the strength of selection due to reproductive interference) increases with the strength of female discrimination. However, when female discrimination is strong and exceeds a threshold, females are able to distinguish males from their own species from individuals from species 2, which limits divergence of the trait in species 1 (see Figure A4(b)). Accurate females choice allows a quasi-similarity of the traits of species 1 and of species 2, without implying heterospecific mating, relaxing the divergent selection exerted by reproductive interference.

**Figure A4:**
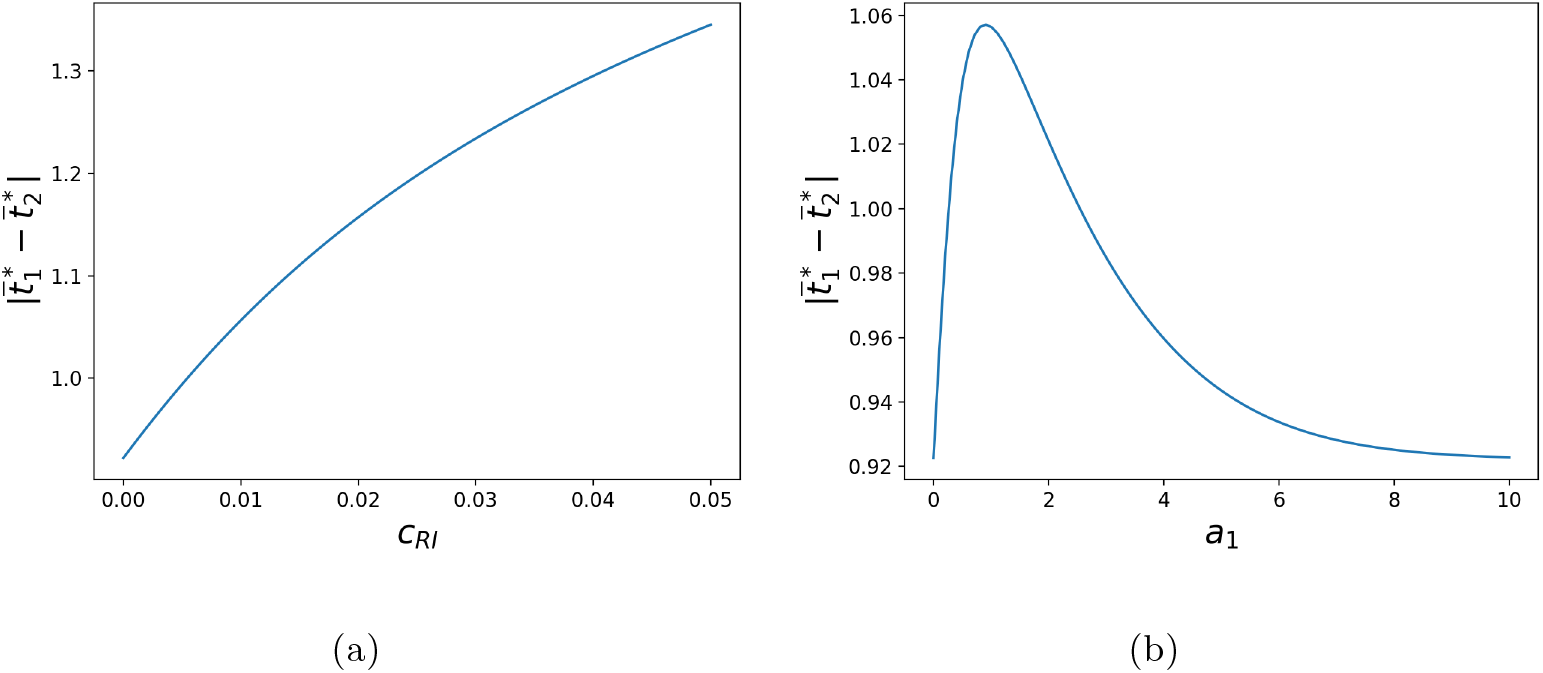
Influence of the strength of (a) reproductive interference *c_RI_* and of (b) female discrimination *a*_1_ on the phenotypic distances between the two species, assuming strong female and predator discriminations. The default parameters values are as follows: *G*_*t*_1__ = *G*_*p*_1__ = 0.01, *a*_1_ = 1, *c_RI_* = 0.01, *b* =1, *d* = 0.02, *λ*_1_ = 0.1, *λ*_2_ = 0.1, *n*_1_ = 10, *n*_2_ = 20, *s*_1_ = 0.0025, *t*_*a*1_ = 0 and *t*_*a*2_ = 1.

##### 5.4.2 Effect of predation

When *a*_1_ and *b* are of order *ε*,

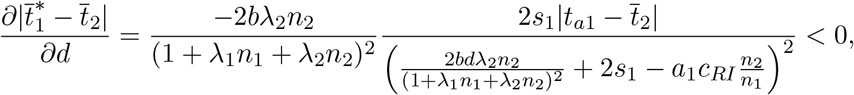

hence the phenotypic distance decreases with the basal predation rate d. Assuming strong female and predator discriminations leads to similar effect (Figure A5(a)). When *a*_1_ and *b* are of order *ε*,

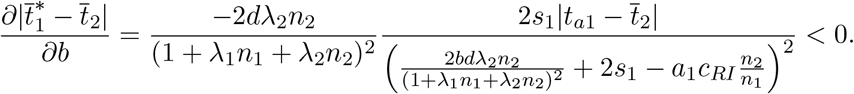

An increase on predator discriminating ability *b* thus also promotes phenotypic convergence as it increases the positive selection on mimicry generated by predation. Surprisingly, when *a*_1_ and *b* are of order 1 and predator discrimination exceeds a certain threshold, increased discrimination no longer promotes accurate mimicry (Figure A5(b)). When *b* is high, individuals from species 1 benefit from protection gained by mimicry only when they look very similar to the individuals of species 2. Reproductive interference makes such high similarity too costly, and therefore divergence of the trait in species 1 becomes promoted (Figure A5(b)).

##### 5.4.3 Effect of the defence level of species 1

Without reproductive interference (*c_RI_* = 0) and when *a*_1_ and *b* are of order *ε*, we have

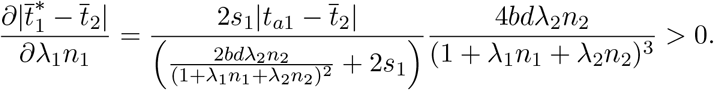

Therefore, the more defended species 1 is, the greater is the phenotypic distance between the two species.

Assuming reproductive interference generates cost (*c_RI_* > 0), the density of species 1 then modulates the fitness cost incurred by reproductive interference. Therefore, we study the effect of the two components of the level of defence of species 1: the individual defence level *λ*_1_ and the density *n*_1_.

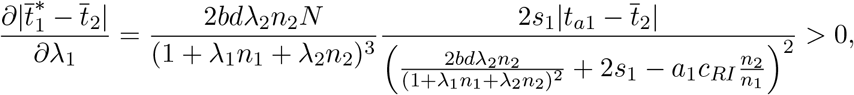

indicates that the individual defence level increases the phenotypic distance. Assuming strong female and predator discriminations leads to similar effect (Figure A6).

**Figure A5:**
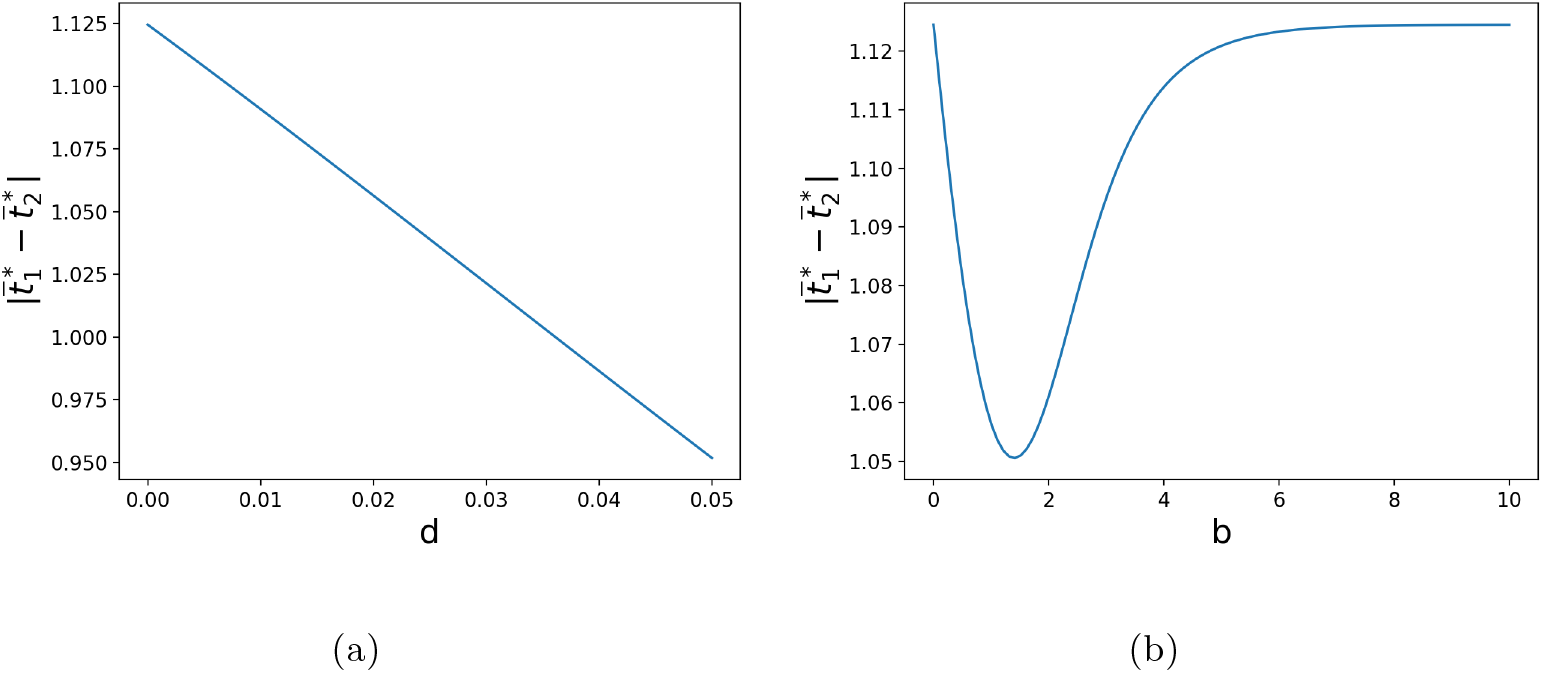
Influence of the strength of predation *d* (a) and of predator discrimination *b* (b) on the phenotypic distances between the two species, assuming strong female and predator discriminations. The default parameters values are as follows: *G*_*t*_1__ = *G*_*p*_1__ = 0.01, *a*_1_ = 1, *c_RI_* = 0.01, *b* =1, *d* = 0.02, *λ*_1_ = 0.1, *λ*_2_ = 0.1, *n*_1_ = 10, *n*_2_ = 20, *s*_1_ = 0.0025, *t*_*a*1_ = 0 and *t*_*a*2_ = 1.

We also have

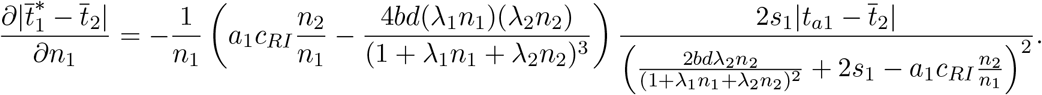

While strong divergence is promoted when species 1 has a low abundance, for larger abundances, the effect of the density on the phenotypic distance between the two species depends on the strength of reproductive interference and predation (Figure A7). When fitness cost due to reproductive interference is great compared to fitness cost due to predation, an increase in the density of species 1 decreases the phenotypic distance, because its reduces the fitness cost due to reproductive interference (similar effect observed assuming strong female and predator discriminations (Figure A7(a))). By contrast, when fitness cost due to reproductive interference is low compared to fitness cost due to predation, an increase in the density of species 1 increases the phenotypic distance because species 1 is better defended (similar effect observed assuming strong female and predator discriminations (see right part of Figure A7(b))).

**Figure A6:**
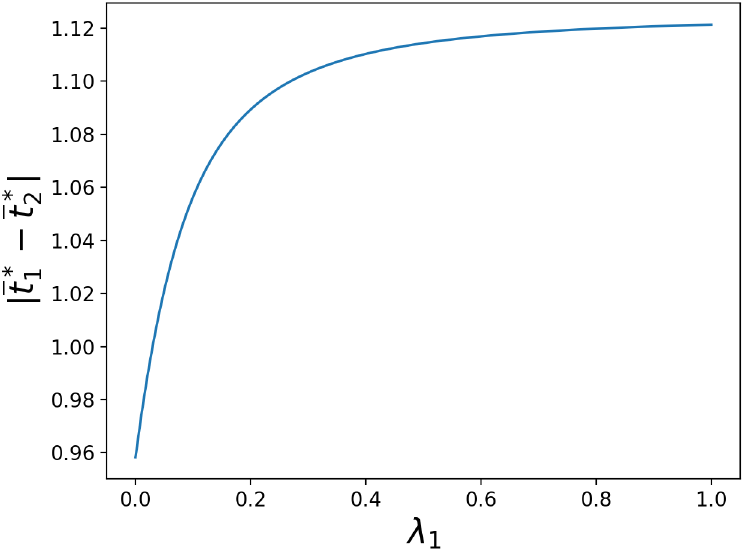
Influence of the individual defence level *λ*_1_ on the phenotypic distances between the two species, assuming strong female and predator discriminations. The default parameters values are as follows: *G*_*t*_1__ = *G*_*p*_1__ = 0.01, *a*_1_ = 1, *c_RI_* = 0.01, *b* =1, *d* = 0.02, *λ*_1_ = 0.1, *λ*_2_ = 0.1, *n*_1_ = 10, *n*_2_ = 20, *s*_1_ = 0.0025, *t*_*a*1_ = 0 and *t*_*a*2_ = 1.

**Figure A7:**
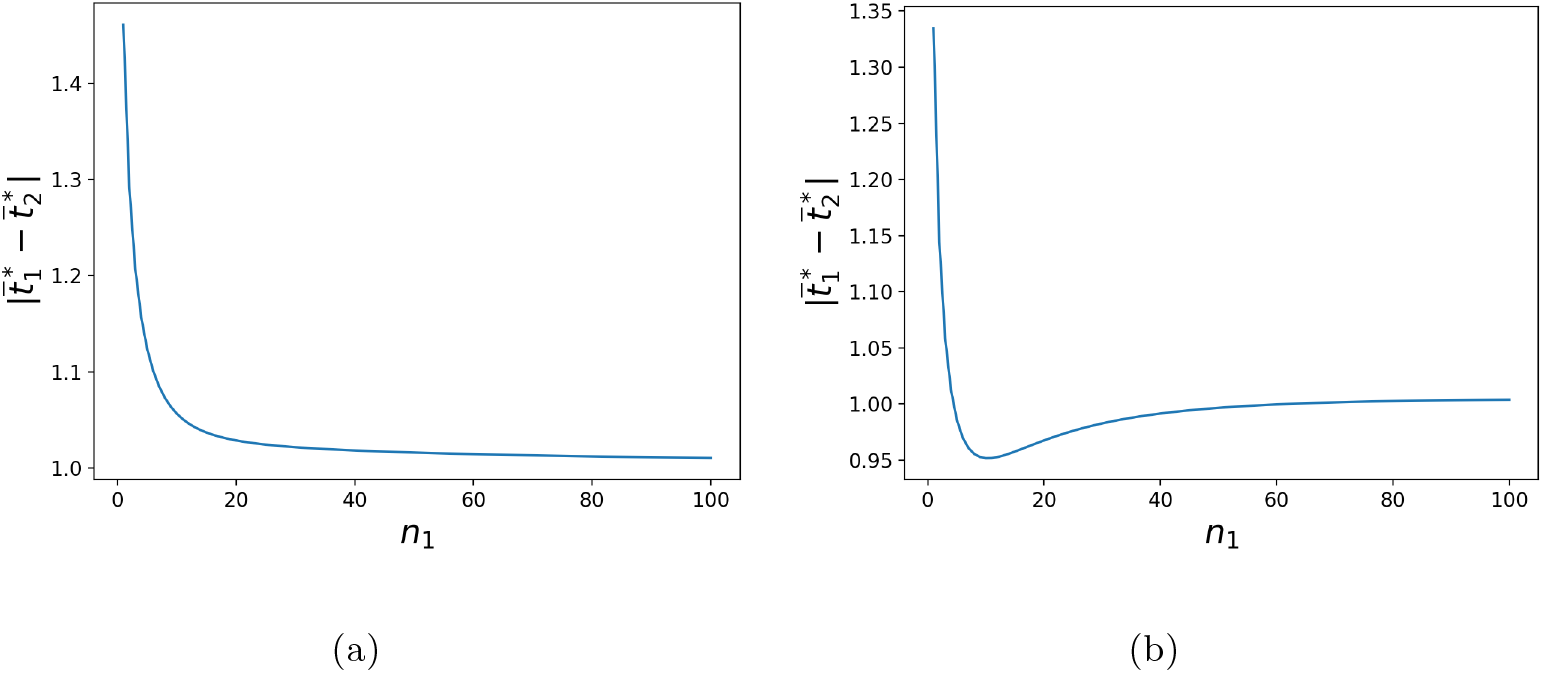
Influence of the density of species 1 *n*_1_ on the phenotypic distances between the two species, when (b) predation is low compared to reproductive interference and (b) when predation is high compared to reproductive interference, assuming strong female and predator discriminations. Simulations were run assuming: (a) *d* = 0.02 or (b) *d* = 0.05. The default parameters values are as follows: *G*_*t*_1__ = *G*_*p*_1__ = 0.01, *a*_1_ = 1, *c_RI_* = 0.01, *b* =1, *λ*_1_ = 0.1, *λ*_2_ = 0.1, *n*_1_ = 10, *n*_2_ = 20, *s*_1_ = 0.0025, *t*_*a*1_ = 0 and *t*_*a*2_ = 1.

##### 5.4.4 Effect of historical constraints promoting the ancestral trait value *t*_*a*1_

When *a*_1_ and *b* are of order *ε*,

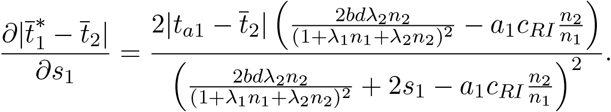

Thus when selection due to predation is stronger than selection due to reproductive interference (i.e. 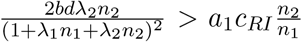), historical constraints promotes the divergence of trait between the two species (similar effect observed assuming strong female and predator discriminations (Figure A8(b))). By contrast, when the selection due to reproductive interference is stronger than selection due to predation (i.e. 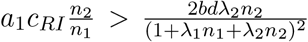), historical constraints limits the trait divergence due to reproductive interference (similar effect observed assuming strong female and predator discriminations (Figure A8(a))).

**Figure A8:**
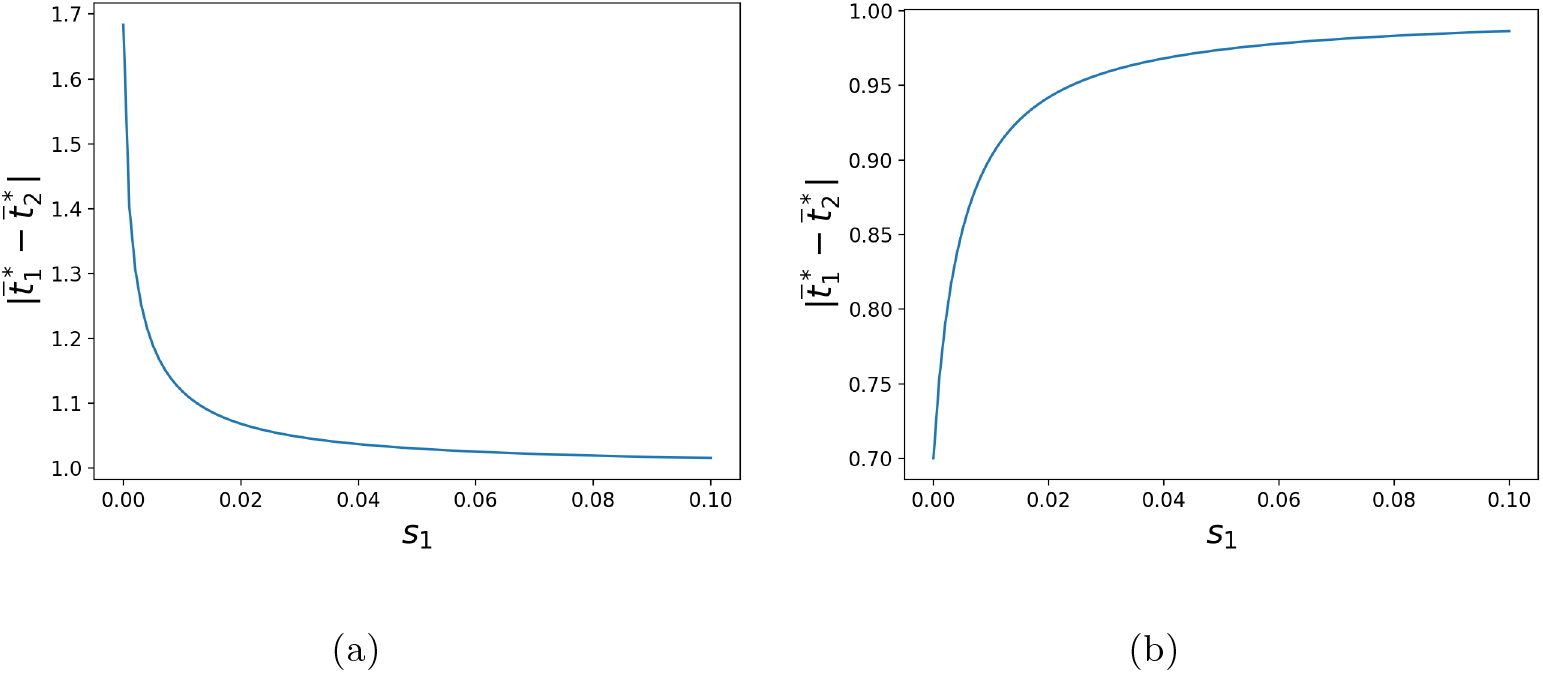
Influence of the strength of selective constraints *s*_1_ on the phenotypic distances between the two specie, when (b) predation is low compared to reproductive interference and (b) when predation is great compared to reproductive interference, assuming strong female and predator discriminations. Simulations were run assuming: (a) *d* = 0.02 or (b) *d* = 0.05. The default parameters values are as follows: *G*_*t*_1__ = *G*_*p*_1__ = 0.01, *a*_1_ = 1, *c_RI_* = 0.01, *b* = 1, *λ*_1_ = 0.1, *λ*_2_ = 0.1, *n*_1_ = 10, *n*_2_ = 20, *s*_1_ = 0.0025, *t*_*a*1_ = 0 and *t*_*a*2_ = 1.

##### 5.4.5 Effect of the defence level of species 2

Finally, we investigate the impact of the composition of species 2 (density *n*_2_ and defense level *λ*_2_) on trait divergence. When *a*_1_ and *b* are of order *ε*, we have:

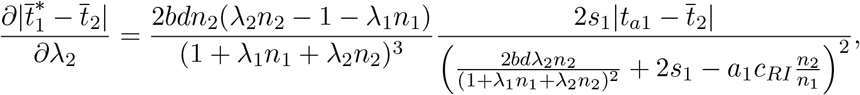

and

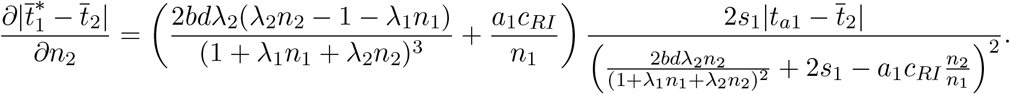

Thus, when the defence level of species 2 (*λ*_2_*n*_2_) is lower than the defence level of species 1 plus one (*λ*_2_*n*_2_ < 1 + *λ*_1_*n*_1_), an increase in the defence level of individuals of species 2 promotes phenotypic similarity between the two species (similar effect observed assuming strong but moderate female and predator discriminations (Figure A9(a))). Indeed, an increase of *λ*_2_*n*_2_ increases the advantage of looking similar to individuals of species 2. More surprisingly, when the defence level of species 2 is greater (*λ*_2_*n*_2_ > 1 + *λ*_1_*n*_1_), the phenotypic distance increases with *λ*_2_*n*_2_ (similar effect observed assuming strong but moderate female and predator discriminations (Figure A9(a))). Under weak (*b* = *O*(*ε*)) or strong but moderate (*e.g. b* = 1) predator discrimination, predators tend to generalize mimetic trait. Then individuals of species 1 gain at least a little protection from species 2, even when displaying a imperfectly mimetic trait. When the level of protection of species 2 is high, the fitness costs linked to predation is low compared to the reproductive interference and decreases with the defence level of individuals of species 2. This decrease in selection pressure generated by predation decreases the phenotype similarity between the two species. When predator discrimination is strong and high (*e.g*. *b* = 10), there is no generalization of imperfect mimics by predators, this effect is no longer observed (see Figure A9(b)).

The density *n*_2_ and the individual defence level of species 2 *λ*_2_ have the same effect on the overall defence level of species 2 (function of the parameter *λ*_2_*n*_2_), and therefore have a similar effect on the phenotypic distance. However, the density of species 2 *n*_2_ also plays a role on the fitness costs generated by reproductive interference. When the strength of reproductive interference is high as compared to predation, the phenotypic distance increases with the density of species 2 (*n*_2_), because it increases the cost of reproductive interference (similar effect observed assuming strong female and predator discriminations (Figure A10(a))). When the strength of reproductive interference is low compared to predation, the density of species 2 (*n*_2_) has a similar effect than the defence level of individuals of species 2 *λ*_2_ on the phenotypic distance (similar effect observed assuming strong but moderate female and predator discriminations (Figure A10(b))).

**Figure A9:**
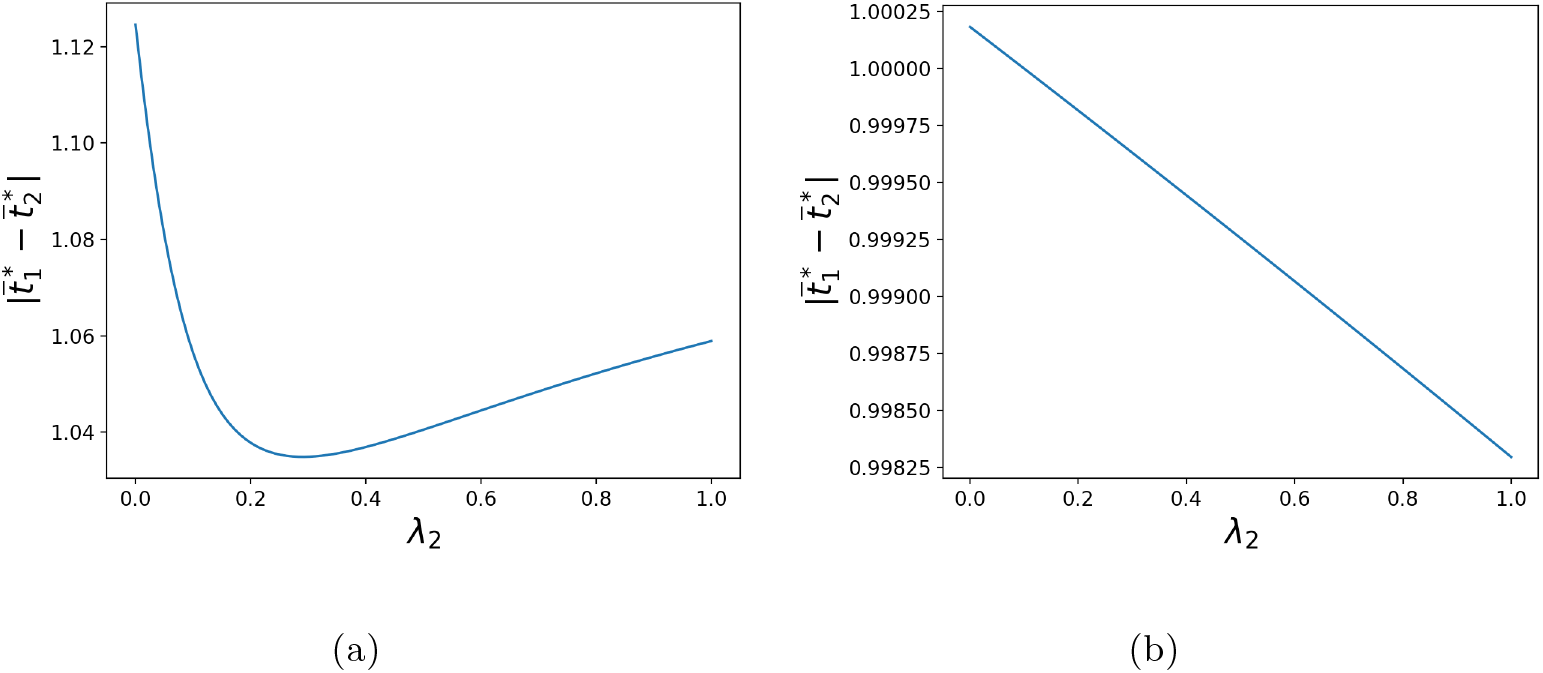
Influence of the mean defence level of individuals of species 2 *λ*_2_ on the phenotypic distances between the two species for (a) moderate (*b* =1) and (b) high (*b* = 10) predator discrimination, assuming strong female and predator discriminations (*i.e*. *a*_1_ = *O*(1) and *b* = *O*(1)). Simulations were run assuming: (a) *a*_1_ = 1 and (b) *a* = 10. The default parameters values are as follows: *G*_*t*_1__ = *G*_*p*_1__ = 0.01, *a*_1_ = 1, *c_RI_* = 0.01, *d* = 0.02, *λ*_1_ = 0.1, *λ*_2_ = 0.1, *n*_1_ = 10, *n*_2_ = 20, *s*_1_ = 0.0025, *t*_*a*1_ = 0 and *t*_*a*2_ = 1.

**Figure A10:**
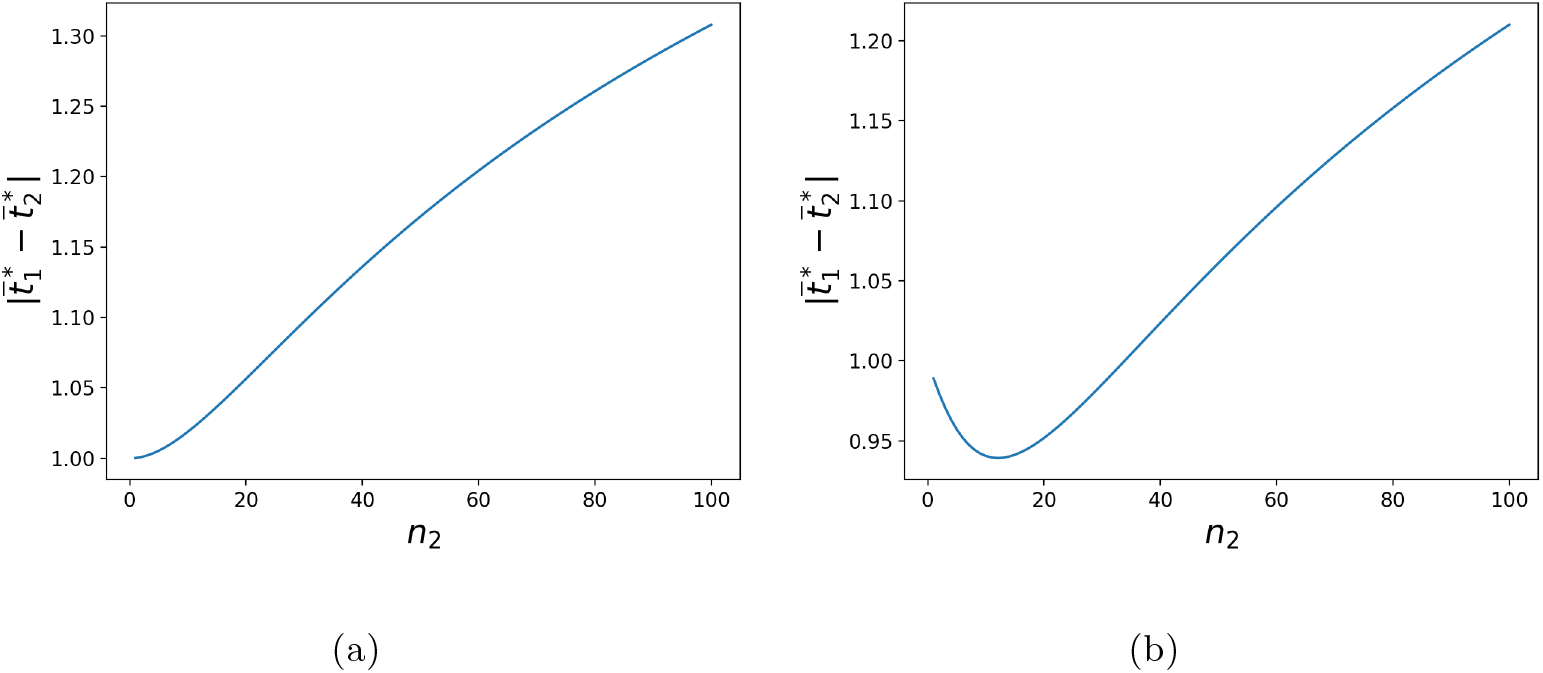
Influence of the density of species 2 *n*_2_ on the phenotypic distances between the two species, when (b) predation is low compared to reproductive interference and (b) when predation is high compared to reproductive interference, assuming strong female and predator discriminations. Simulations were run assuming: (a) *d* = 0.02 or (b) *d* = 0.05. The default parameters values are as follows: *G*_*t*_1__ = *G*_*p*_1__ = 0.01, *a*_1_ = 1, *c_RI_* = 0.01, *b* =1, *λ*_1_ = 0.1, *λ*_2_ = 0.1, *n*_1_ = 10, *n*_2_ = 20, *s*_1_ = 0.0025, *t*_*a*1_ = 0 and *t*_*a*2_ = 1.

### 6 Evolution of mimicry between two interacting species

In this section we focus on the general case where traits and preferences5 co-evolve in the two species. In Section 5.1 we compute the values of trait and preference that cancel the leading term of the selection vectors in both species. In Section 5.2 we study when trait and preference converge toward the point found in the previous section. In Section 5.3 we detail the impact of each parameter of the interspecific phenotypic distance. In Section 5.4 we study the impact of historical constraints on phenotypic divergence. In section 5.5 we study the impact of predator discrimination without reproductive interference. In section 5.5 we study the impact of the relative defence level on traits co-evolution.

#### 6.1 Quasi equilibria

We search mean values of traits and preferences at equilibrium 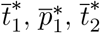 and 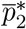 such as the leading terms of 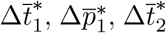 and 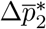 are null, or equivalently, 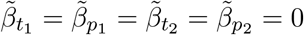 with

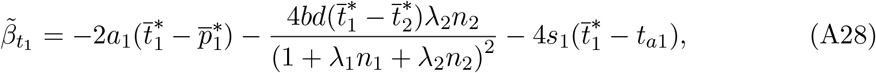

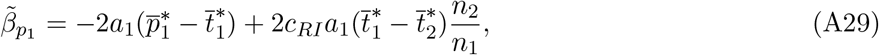

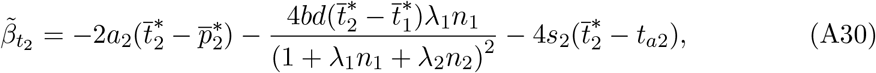

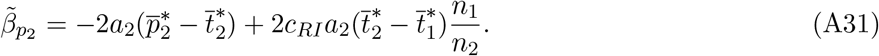

##### Lemma 3.

*Let us introduce for i* ≠ *j* ∈ {1, 2}^2^,

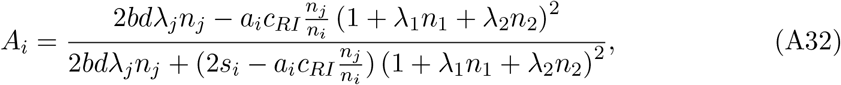

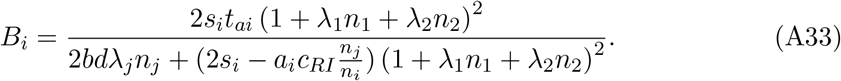

*If* 2*bdλ_j_n_j_* + (2*s_i_* – *a_i_c_RI_*(*n_j_/n_i_*)) (1 + *λ*_1_*n*_1_ + *λ*_2_*n*_2_)^2^ = 0 *for i* ∈ {1, 2} *and if A*_1_*A*_2_ ≠ 1 *there is a unique quasi equilibrium point given for i* ∈ {1, 2} *by*

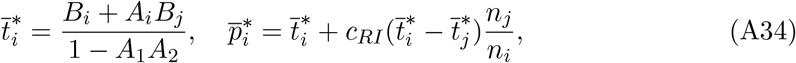

**Remark.** We do not give the quasi equilibrium traits and preferences when one of the following conditions is verified:

- 2*bdλ*_2_*n*_2_ + (2*s*_1_ – *a*_1_*c_RI_*(*n*_2_/*n*_1_)) (1 + *λ*_1_*n*_1_ + *λ*_2_*n*_2_)^2^ = 0,
- 2*bdλ*_1_*n*_1_ + (2*s*_2_ – *a*_2_*c_RI_*(*n*_1_/*n*_2_)) (1 + *λ*_1_*n*_1_ + *λ*_2_*n*_2_)^2^ = 0,
- *A*_1_*A*_2_ = 1.

Indeed investigating those cases leads to a long enumeration of subcases with different quasi equilibrium points, and those cases are negligible compared with all the possible combinations of parameters. We thus choose to not give this enumeration.

*Proof*. Let 2*bdλ_j_n_j_* + (2*s_i_* – *a_i_c_RI_*(*n_j_/n_i_*)) (1 + *λ*_1_*n*_1_ + *λ*_2_*n*_2_)^2^ ≠ 0 for *i* ≠ *j* ∈ {1, 2}^2^. Using the results of the first model (see Appendix 1.3) we have

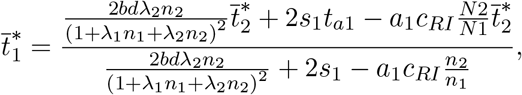

and

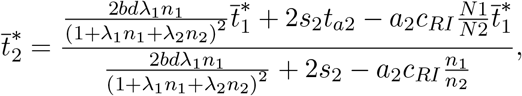

which can be rewritten

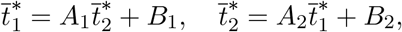

where (*A_i_,B_i_*), *i* ∈ {1, 2} have been defined in (A32)-(A33).

If *A*_1_*A*_2_ ≠ 1 there are unique values for traits at quasi equilibrium given by 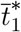 and 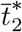 defined in (A34). Using previous results we have the associated mean preferences at quasi equilibrium 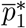 and 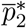 defined in (A34).

#### 6.2 Fast and slow dynamics

In this section we study when trait and preference in the both species converge towards the quasi equilibrium point.

##### Lemma 4.

*Let us introduce*

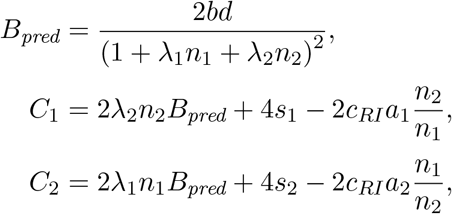

*and recall the definitions of A*_1_ *and A*_2_ in (A32).

*1. If C*_1_ > 0, *C*_2_ > 0 *and* 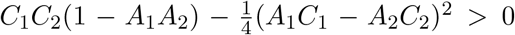 *the quantities* 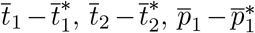 *and* 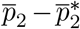 *become of order* 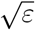 *when the number of generation goes to infinity, with* 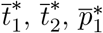 *and* 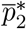 *defined by* (A34).

*2. If C*_1_ < 0, *C*_2_ < 0 *and* 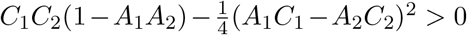, *traits and preferences become very large*.

**Remark.** When 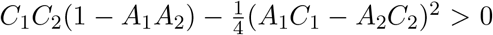 or *C*_1_*C*_2_ < 0 we are not able to analytically determine the convergence or the divergence of traits and preferences. To infer such information we simulate the dynamic of traits and preferences considering the leading term of the selection coefficients.

*Proof*. As previously, we can decompose the dynamics into two steps. In the first one the leading order terms of the selection coefficient are the terms describing sexual selection and cost of choosiness. Using computations similar to the ones in the previous model we can approximate 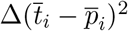 for *i* ∈ {1, 2} by

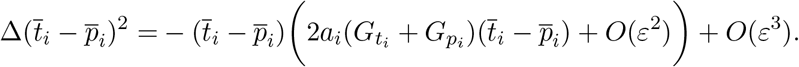

Using the same reasoning that in the previous model we may prove that above a certain number of generations *t*_1_ – *p*_1_ and *t*_2_ – *p*_2_ are of order *ε*.

We now look at the variation of

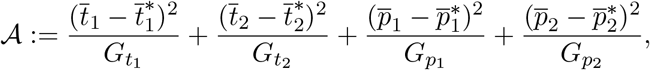

with 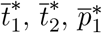 and 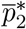 defined by (A34).

By definition,

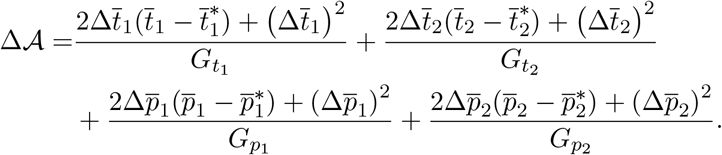

When 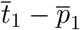 and 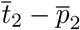 are of order *ε*, 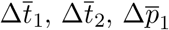 and 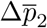 are of order *ε*^2^. We then neglect the terms proportional to 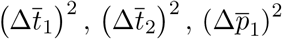 or 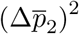:

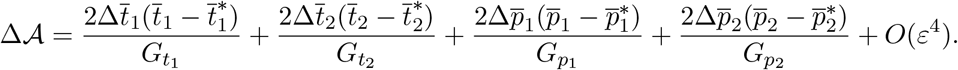

By replacing the terms 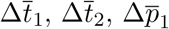 and 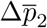 using (12) and simplifying, we obtain:

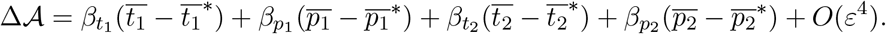

Using that for *i* ∈ {1, 2}, 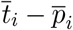 and 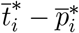 are of order *ε*, we obtain similarly as in (A22)

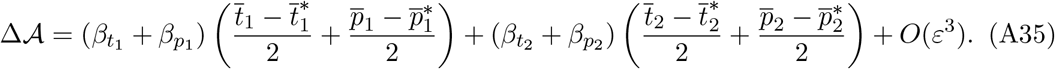

Using (A3) and (A4) and simplifying, we obtain:

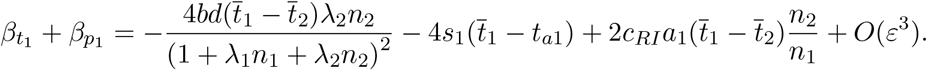

By substracting 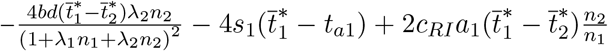 which is equal to zero (because 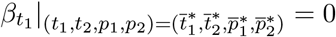 and 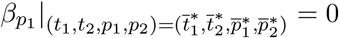, we get:

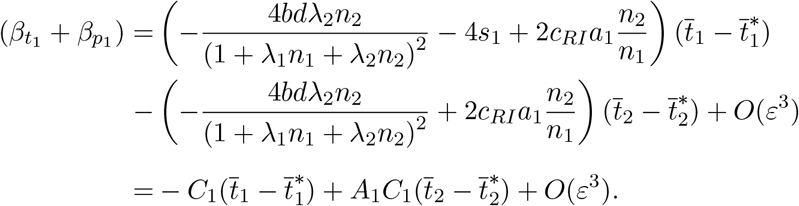

Because for *i* ∈ {1, 2}, 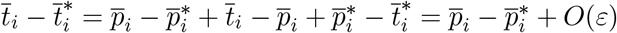 we also have

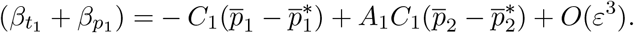

We thus get

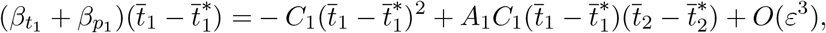

and

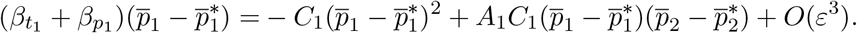

Hence using (A35), the symmetry of the model between populations 1 and 2, and that for *i* ∈ {1, 2}, 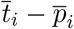 and 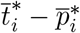 are of order *ε*, we obtain:

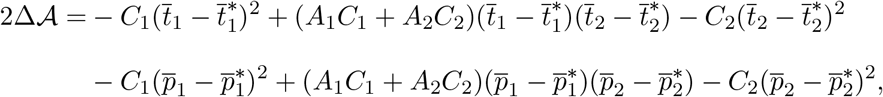

with can be rewritten in matrix form

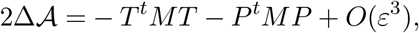

with

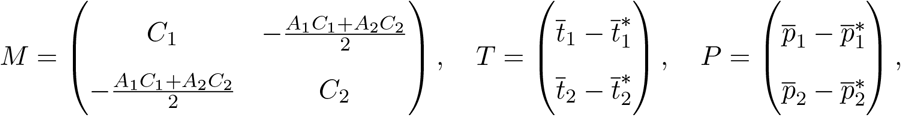

and *T^t^* and *P^t^* being the transposed of *T* and *P* respectively.

**First case:** *C*_1_ > 0, *C*_2_ > 0 **and** 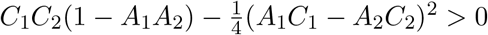.

According to the Sylvester’s criterion the matrix *M* is positive-definite. Using computations similar to the ones derived to prove (A17), we may prove that after a number of generations large enough, 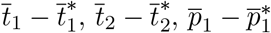 and 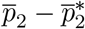 become of order *ε*^1/2^.

**Second case:** *C*_1_ < 0, *C*_2_ < 0 **and** 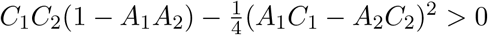.

According to the Sylvester’s criterion the matrix – *M* is positive-definite. Therefore 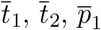 and 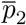 become very large.

#### 6.3 Effect of different parameters on the level of trait divergence 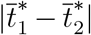 assuming weak or strong female and predator discriminations

We want to verify that the different parameters have similar effect on phenotypic distance between species in this model than in the first. To this aim we plot the effect of different parameters on the level of trait divergence assuming whether (using the analytical approximation when possible or numerical simulations) or (using numerical simulations). The following plots confirm that the parameters have a similar effect on the phenotypic distance.

**Figure A11:**
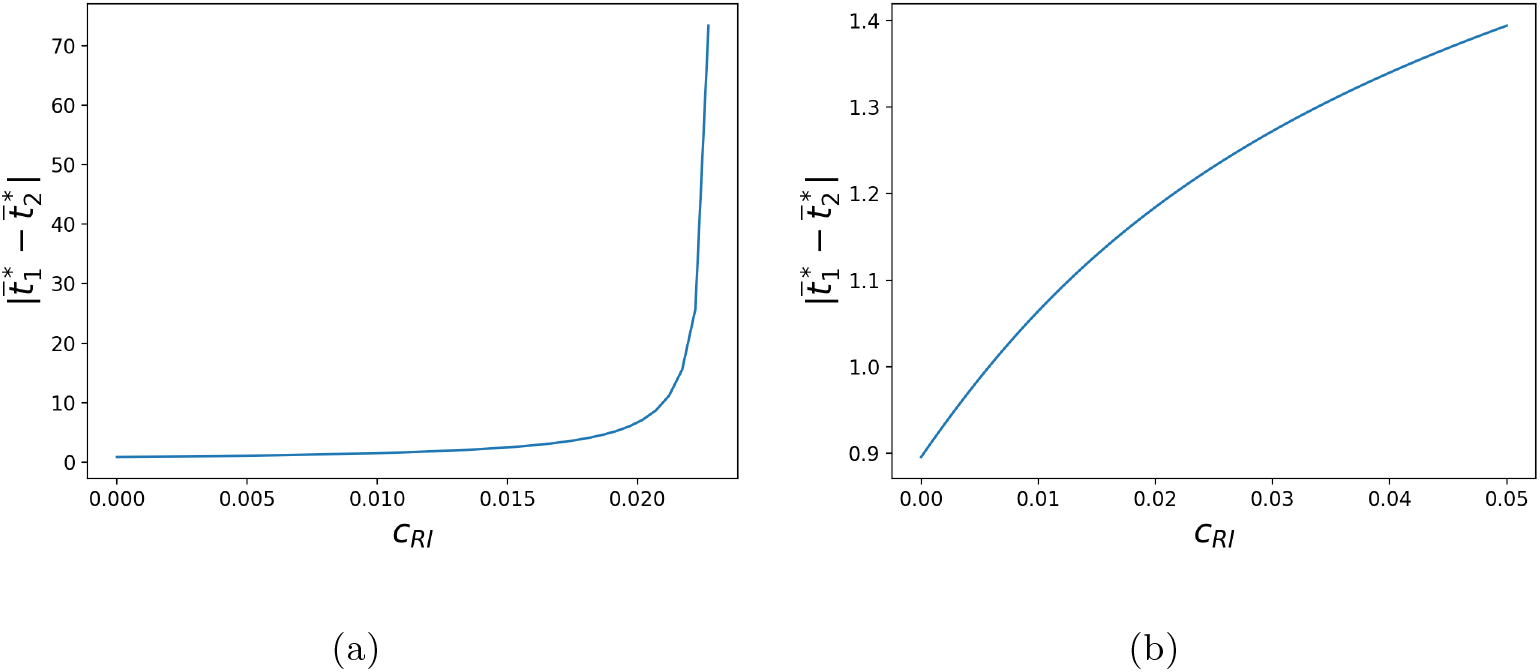
Influence of the strength of reproductive interference *c_RI_* on the phenotypic distances between the two species, assuming (a) weak or (b) strong female and predator discriminations. Simulations were run assuming (a) *a*_1_ = *a*_2_ = *b* = 0.01 or (b) *a*_1_ = *a*_2_ = *b* =1. The default parameters values are as follows: *G*_*t*_1__ = *G*_*p*_1__ = *G*_*t*_2__ = *G*_*p*_2__ = 0.01, *d* = 0.02, *λ*_1_ = 0.1, *λ*_2_ = 0.1, *n*_1_ = 10, *n*_2_ = 20, *s*_1_ = 0.0000025, *s*_2_ = 0.0000025, *t*_*a*1_ = 0, *t*_*a*2_ = 1.

**Figure A12:**
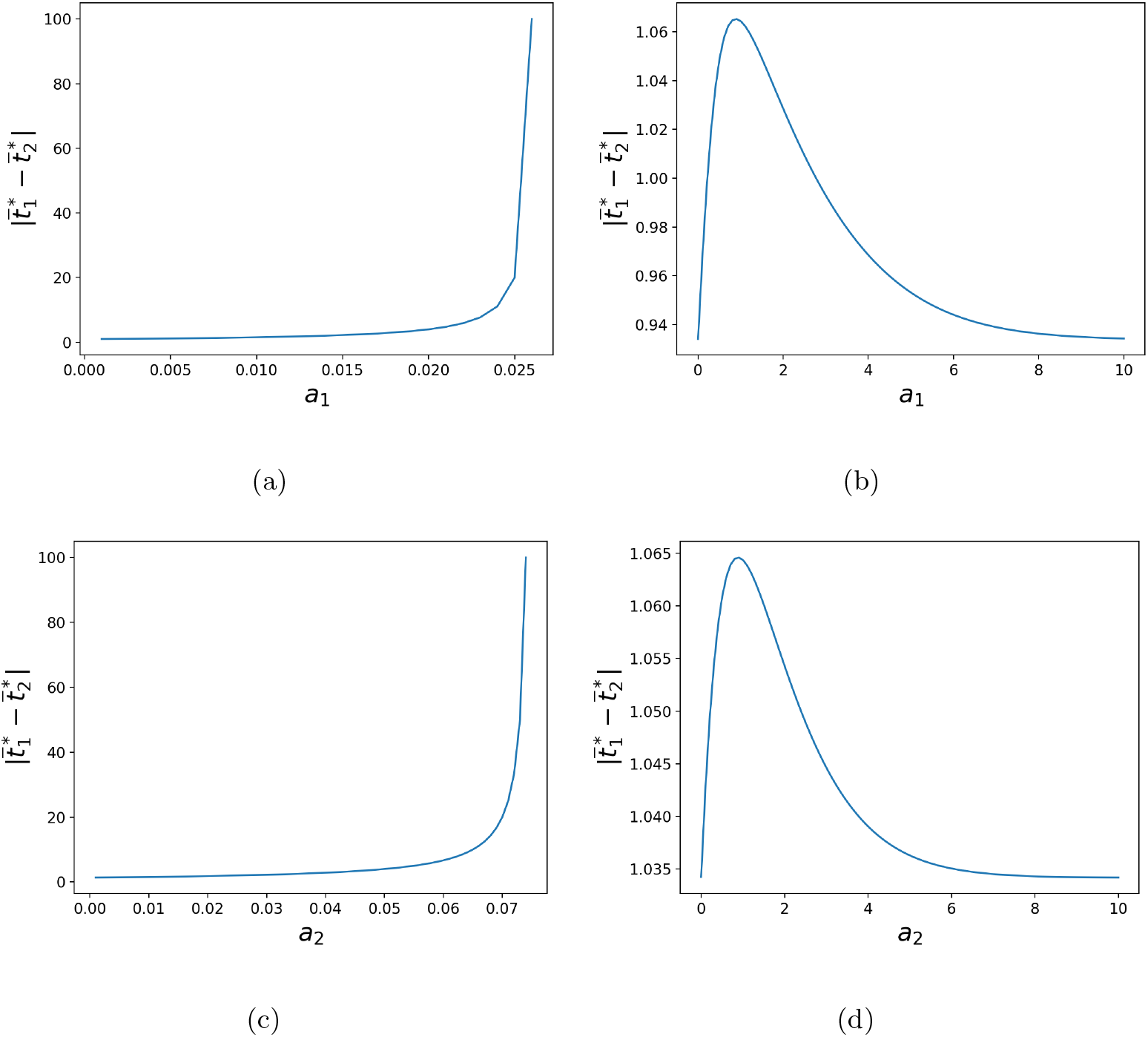
Influence of female discrimination in species (a)(b) 1 *a*_1_ and (c)(d) 2 *a*_1_ on the phenotypic distances between the two species, assuming (a)(c) weak or (b)(d) strong female and predator discriminations. Simulations were run assuming (a) *a*_2_ = *b* = 0.01, (b) *a*_2_ = *b* = 1, (c) *a*_1_ = *b* = 0.01 or (d) *a*_1_ = *b* = 1. The default parameters values are as follows: *G*_*t*_1__ = *G*_*p*_1__ = *G*_*t*_2__ = *G*_*p*_2__ = 0.01, *c_RI_* = 0.01, *d* = 0.02, *λ*_1_ = 0.1, *λ*_2_ = 0.1, *n*_1_ = 10, *n*_2_ = 20, *s*_1_ = 0.0000025, *s*_2_ = 0.0000025, *t*_*a*1_ = 0, *t*_*a*2_ = 1.

**Figure A13:**
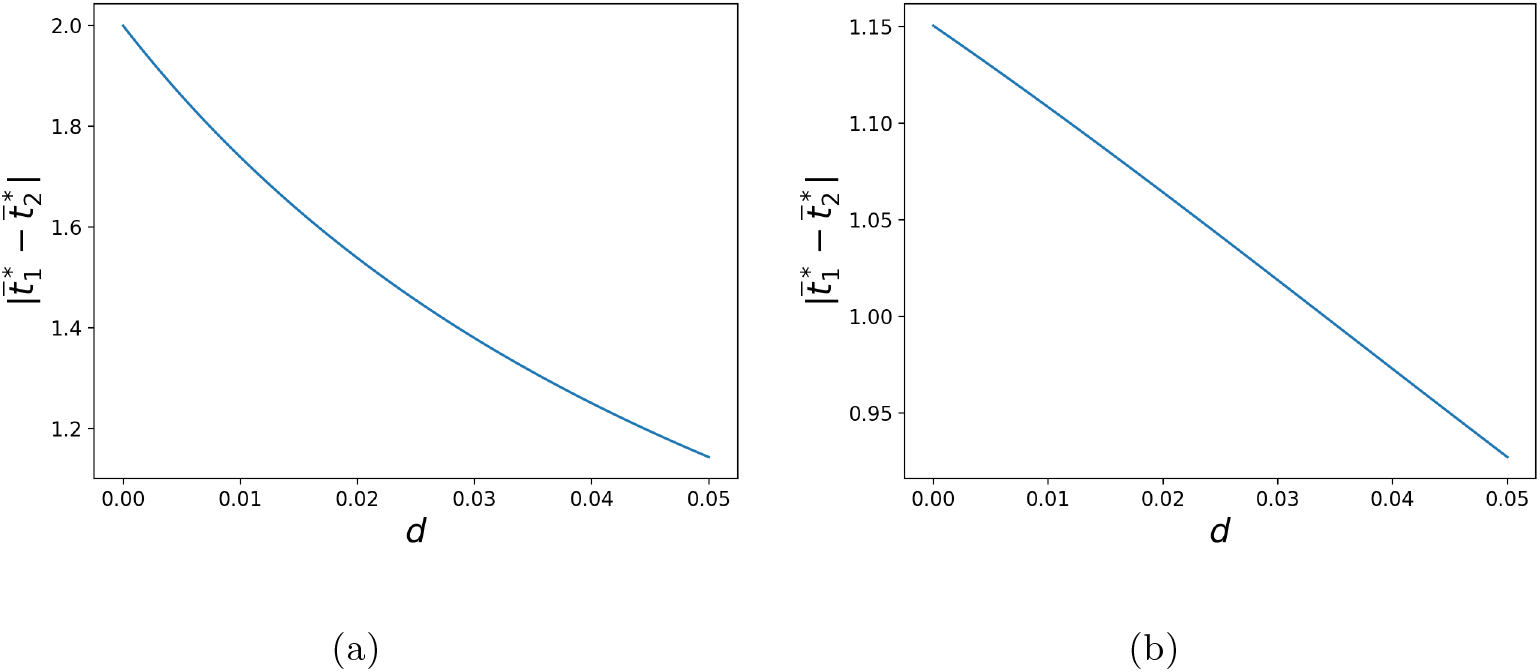
Influence of the strength of predation *d* on the phenotypic distances between the two species, assuming (a) weak or (b) strong female and predator discriminations. Simulations were run assuming (a) *a*_1_ = *a*_2_ = *b* = 0.01, (b) *a*_1_ = *a*_2_ = *b* =1. The default parameters values are as follows: *G*_*t*_1__ = *G*_*p*_1__ = *G*_*t*_2__ = *G*_*p*_2__ = 0.01, *c_RI_* = 0.01, *λ*_1_ = 0.1, *λ*_2_ = 0.1, *n*_1_ = 10, *n*_2_ = 20, *s*_1_ = 0.0000025, *s*_2_ = 0.0000025, *t*_*a*1_ = 0, *t*_*a*2_ = 1.

**Figure A14:**
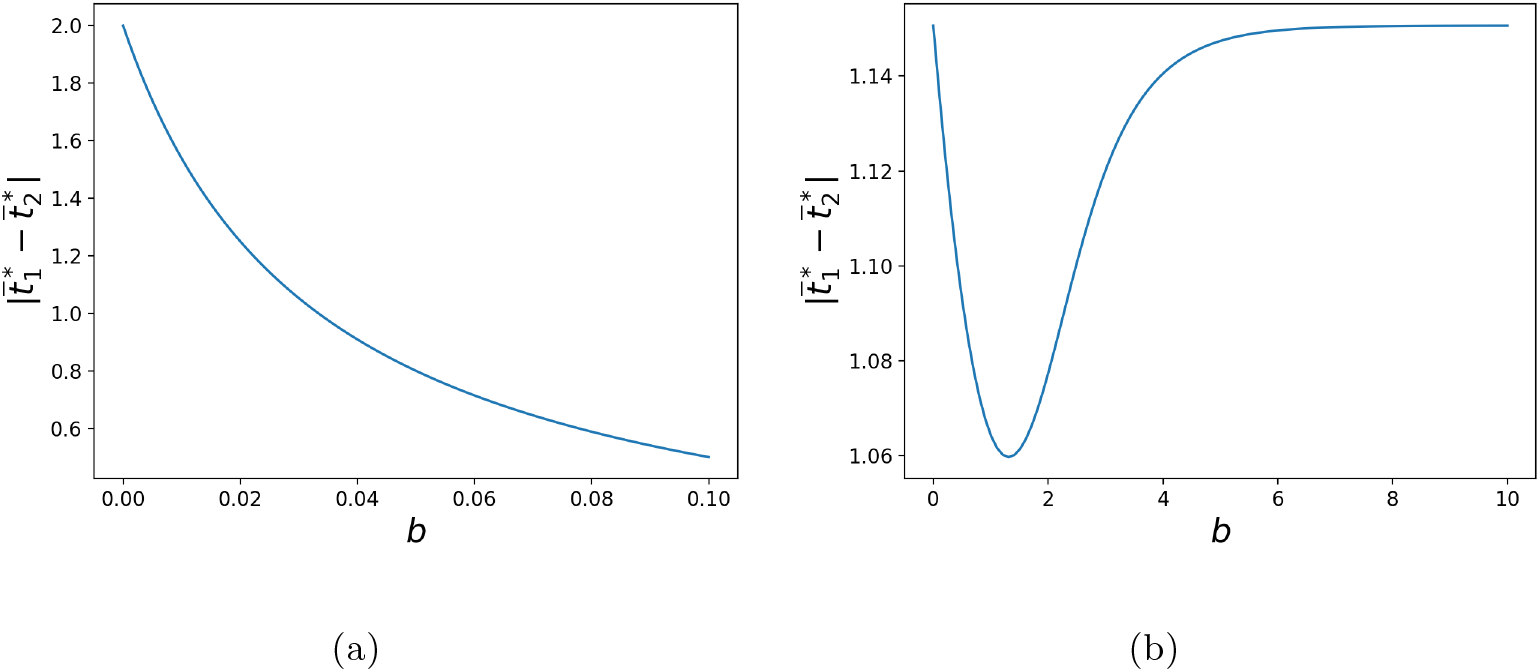
Influence of predator discrimination *b* on the phenotypic distances between the two species, assuming (a) weak or (b) strong female and predator discriminations. Simulations were run assuming (a) *a*_1_ = *a*_2_ = 0.01, (b) *a*_1_ = *a*_2_ = 1. The default parameters values are as follows: *G*_*t*_1__ = *G*_*p*_1__ = *G*_*t*_2__ = *G*_*p*_2__ = 0.01, *c_RI_* = 0.01, *d* = 0.02, *λ*_1_ = 0.1, *λ*_2_ = 0.1, *n*_1_ = 10, *n*_2_ = 20, *s*_1_ = 0.0000025, *s*_2_ = 0.0000025, *t*_*a*1_ = 0, *t*_*a*2_ = 1.

**Figure A15:**
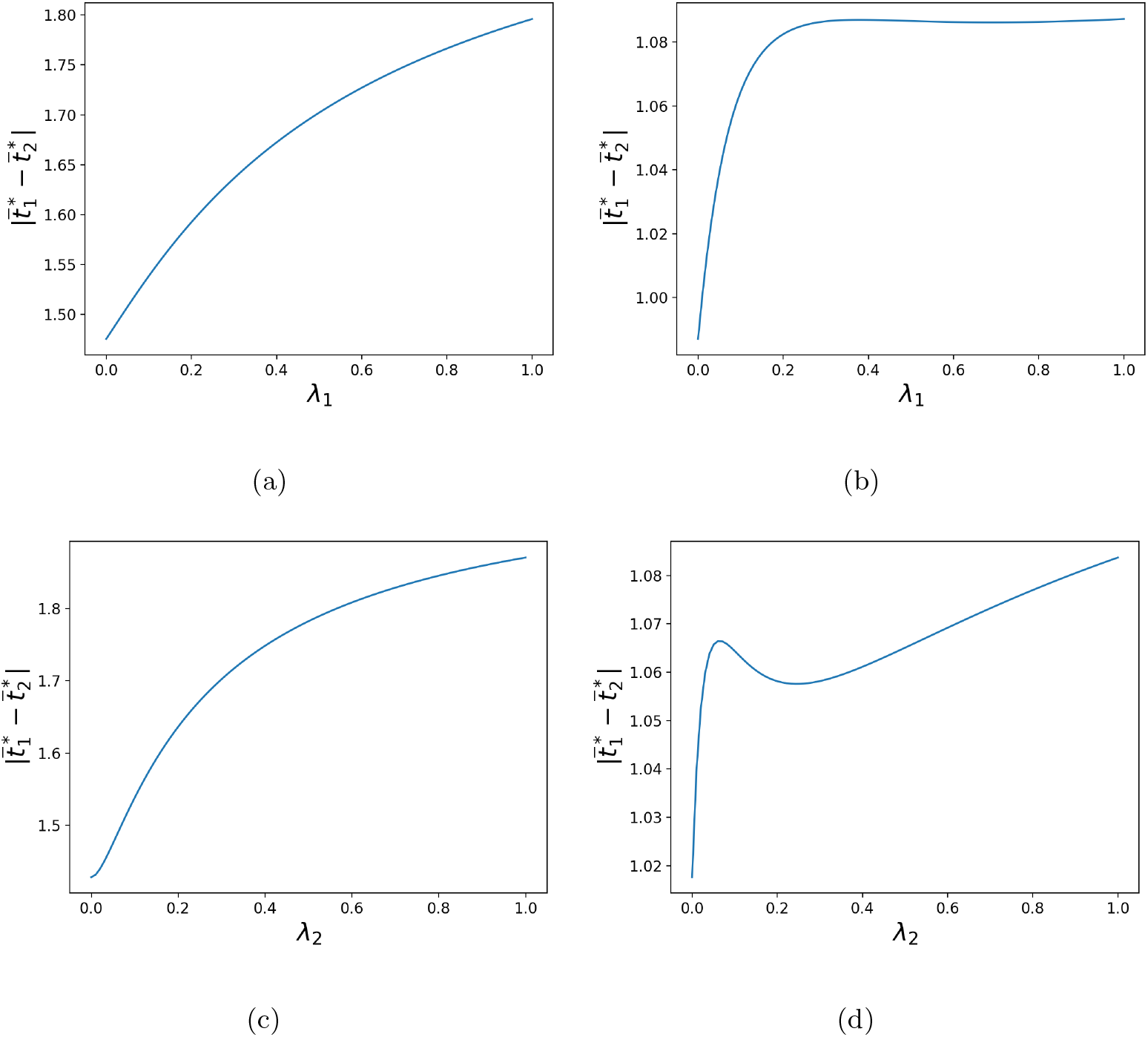
Influence of the defence level of (a)(b) species 1 *λ*_1_ and of (c)(d) species 2 *λ*_2_ on the phenotypic distances between both species, assuming (a)(c) weak or (b)(d) strong female and predator discriminations. Simulations were run assuming (a)(c) *a*_1_ = *a*_2_ = *b* = 0.01, (b)(d) *a*_1_ = *a*_2_ = *b* =1. The default parameters values are as follows: *G*_*t*_1__ = *G*_*p*_1__ = *G*_*t*_2__ = *G*_*p*_2__ = 0.01, *c_RI_* = 0.01, *d* = 0.02, *λ*_1_ = 0.1, *λ*_2_ = 0.1, *n*_1_ = 10, *n*_2_ = 20, *s*_1_ = 0.0000025, *s*_2_ = 0.0000025, *t*_*a*1_ = 0, *t*_*a*2_ = 1.

**Figure A16:**
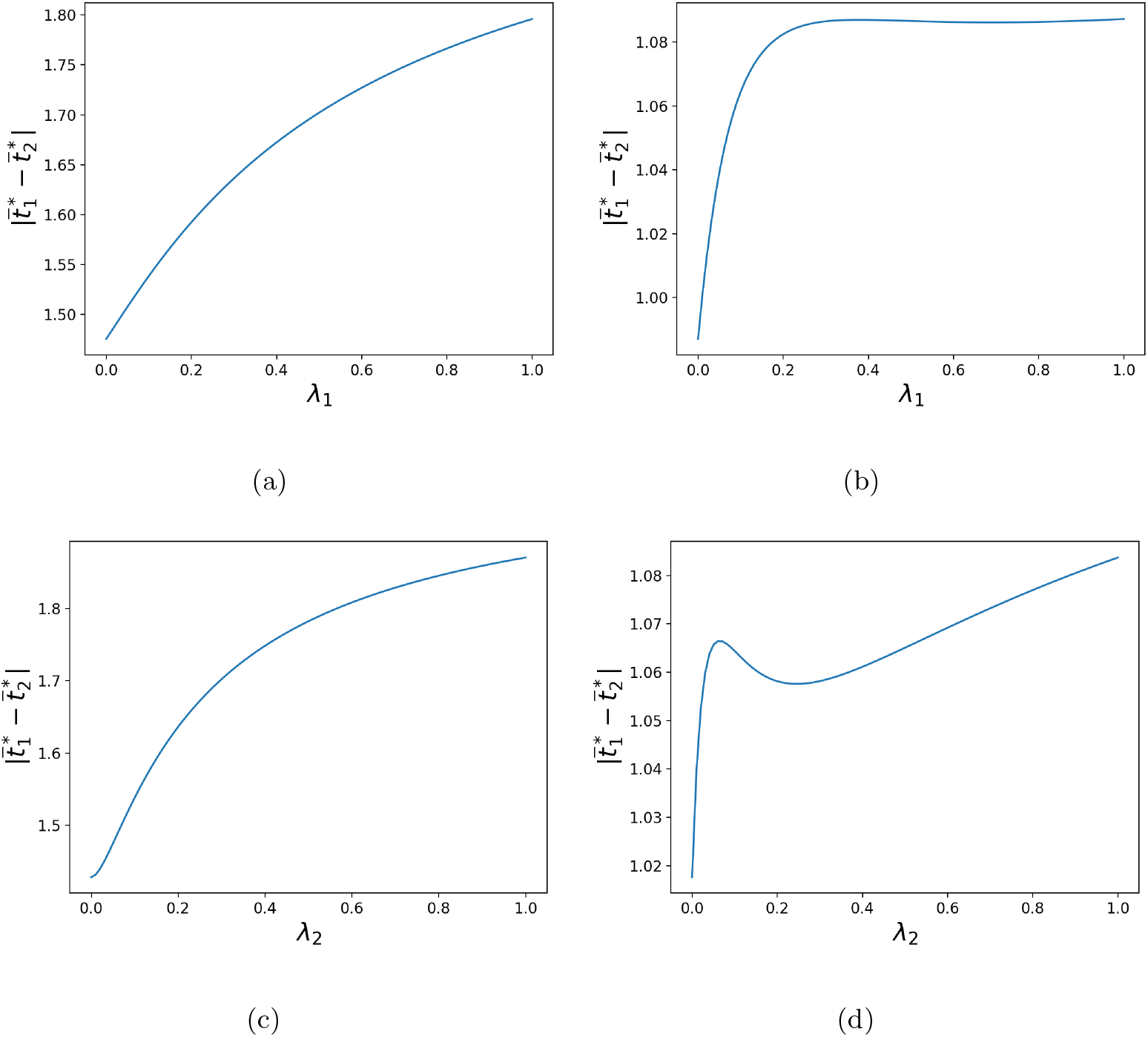
Influence of the defence level of (a)(b) species 1 *λ*_1_ and of (c)(d) species 2 *λ*_2_ on the phenotypic distances between both species, assuming (a)(c) weak or (b)(d) strong female and predator discriminations. Simulations were run assuming (a)(c) *a*_1_ = *a*_2_ = *b* = 0.01, (b)(d) *a*_1_ = *a*_2_ = *b* =1. The default parameters values are as follows: *G*_*t*_1__ = *G*_*p*_1__ = *G*_*t*_2__ = *G*_*p*_2__ = 0.01, *c_RI_* = 0.01, *d* = 0.02, *λ*_1_ = 0.1, *λ*_2_ = 0.1, *n*_1_ = 10, *n*_2_ = 20, *s*_1_ = 0.0000025, *s*_2_ = 0.0000025, *t*_*a*1_ = 0, *t*_*a*2_ = 1.

**Figure A17:**
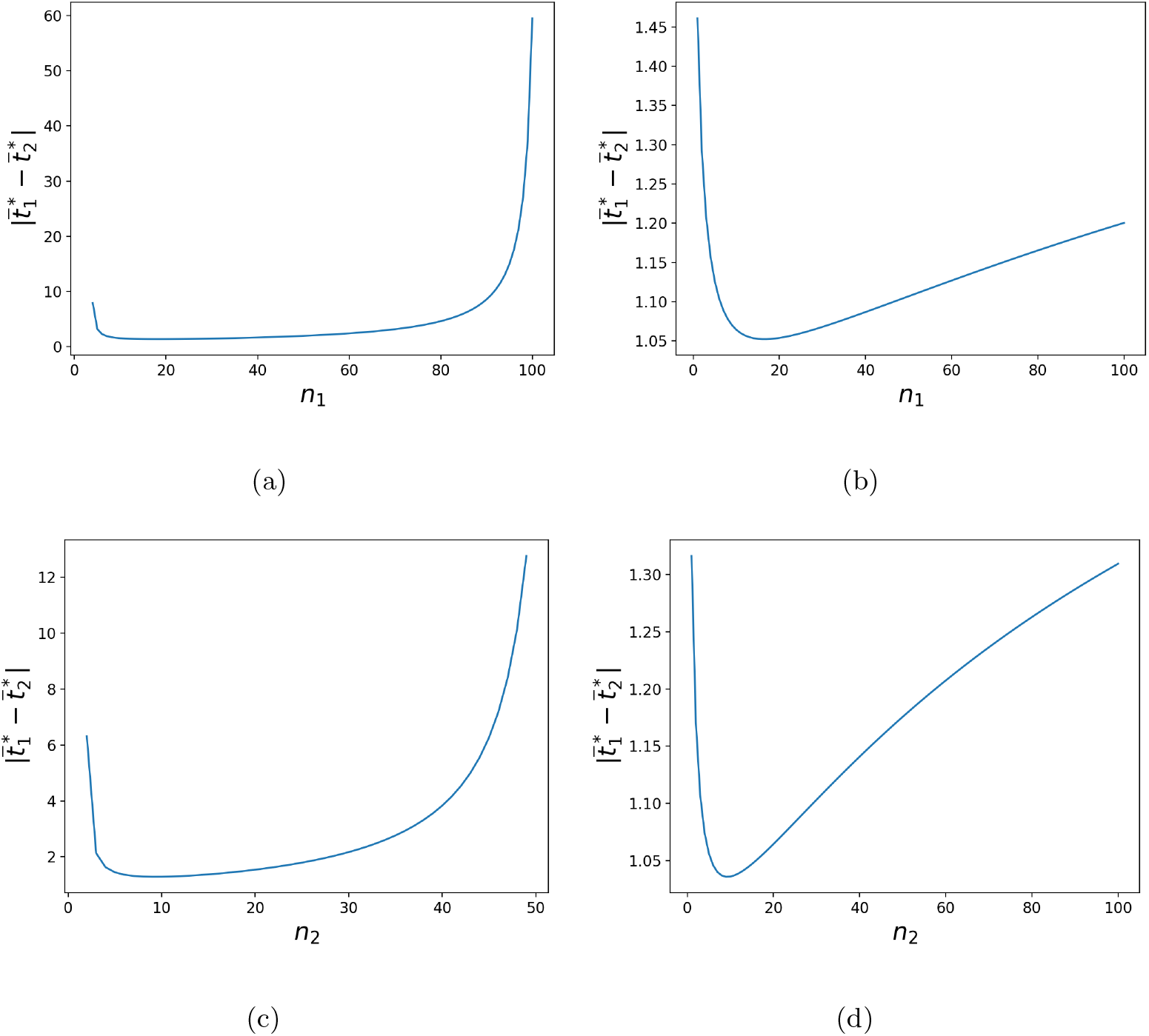
Influence of the density of (a)(b) species 1 *n*_1_ and of (c)(d) species 2 *n*_2_ on the phenotypic distances between the two species, assuming (a)(c) weak or (b)(d) strong female and predator discriminations. Simulations were run assuming (a) *a*_1_ = *a*_2_ = *b* = 0.01, (b) *a*_1_ = *a*_2_ = *b* =1. The default parameters values are as follows: *G*_*t*_1__ = *G*_*p*_1__ = *G*_*t*_2__ = *G*_*p*_2__ = 0.01, *c_RI_* = 0.01, *d* = 0.02, *λ*_1_ = 0.1, *λ*_2_ = 0.1, *n*_1_ = 10, *n*_2_ = 20, *s*_1_ = 0.0000025, *s*_2_ = 0.0000025, *t*_*a*1_ = 0, *t*_*a*2_ = 1.

**Figure A18:**
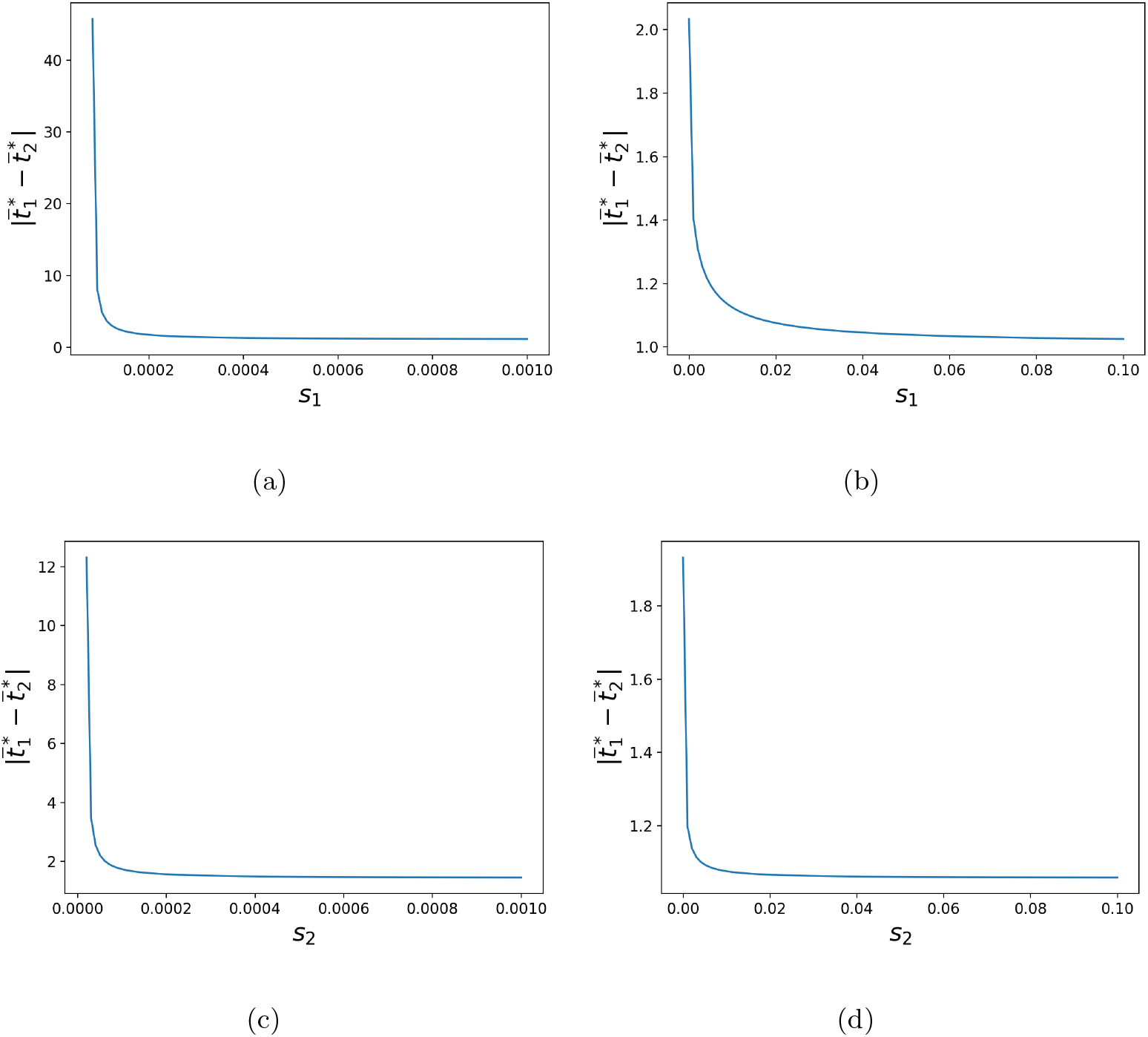
Influence of the strength of selective constraints in (a)(b) species 1 *s*_1_ and in (c)(d) species 2 *s*_2_ on the phenotypic distances between the two species, assuming (a)(c) weak or (b)(d) strong female and predator discriminations. Simulations were run assuming (a)(c) *a*_1_ = *a*_2_ = *b* = 0.01, (b)(d) *a*_1_ = *a*_2_ = *b* =1. The default parameters values are as follows: *G*_*t*_1__ = *G*_*p*_1__ = *G*_*t*_2__ = *G*_*p*_2__ = 0.01, *c_RI_* = 0.01, *d* = 0.02, *λ*_1_ = 0.1, *λ*_2_ = 0.1, *n*_1_ = 10, *n*_2_ = 20, *s*_1_ = 0.0000025, *s*_2_ = 0.0000025, *t*_*a*1_ = 0, *t*_*a*2_ = 1.

**Figure A19:**
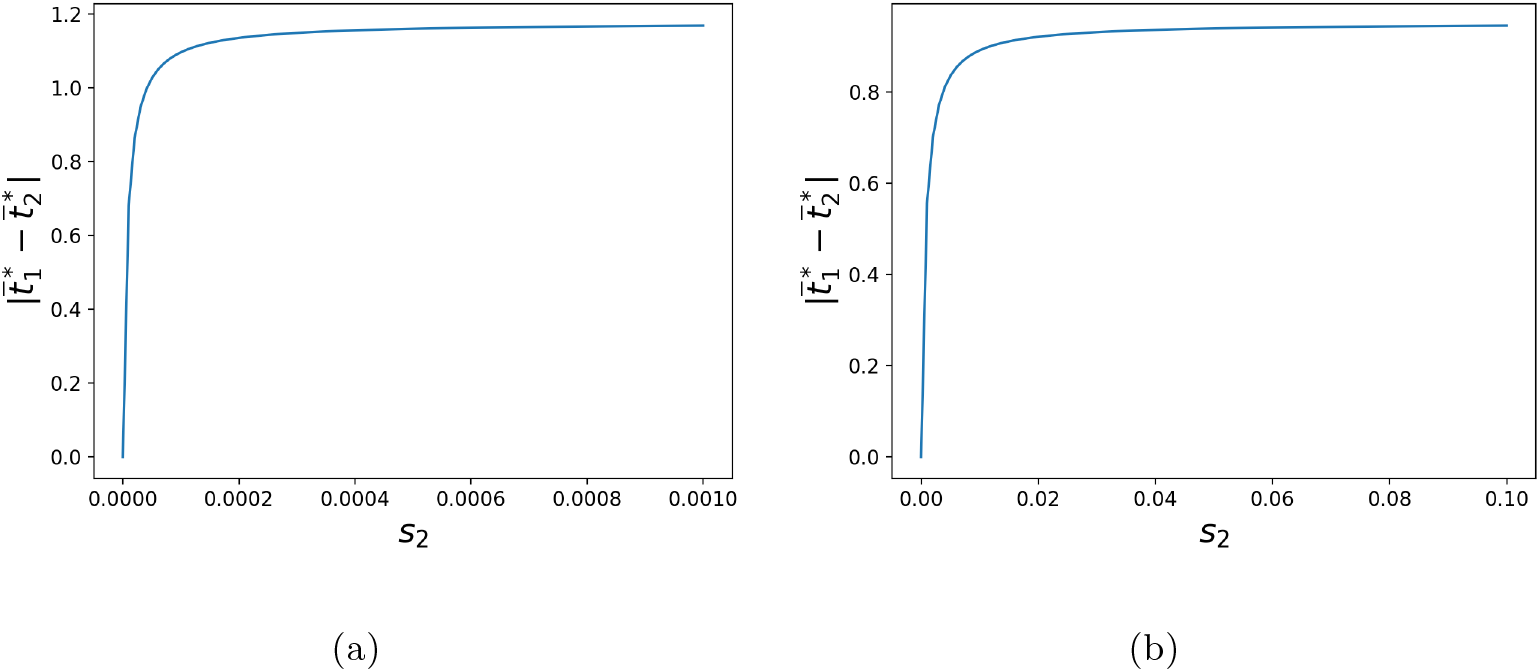
Influence of the strength of selective constraints in species 2 *s*_2_ on the phenotypic distances between the two species, when predation is great compare to reproductive interference, assuming (a) weak or (b) strong female and predator discriminations. Simulations were run assuming (a) *a*_1_ = *a*_2_ = *b* = 0.01, (b) *a*_1_ = *a*_2_ = *b* =1. The default parameters values are as follows: *G*_*t*_1__ = *G*_*p*_1__ = *G*_*t*_2__ = *G*_*p*_2__ = 0.01, *c_RI_* = 0.01, *d* = 0.05, *λ*_1_ = 0.1, *λ*_2_ = 0.1, *n*_1_ = 10, *n*_2_ = 20, *s*_1_ = 0.0000025, *s*_2_ = 0.0000025, *t*_*a*1_ = 0, *t*_*a*2_ = 1.

#### 6.4 Historical constraints promoting ancestral trait values strongly modulate phenotypic divergence driven by reproductive interference

**Figure A20:**
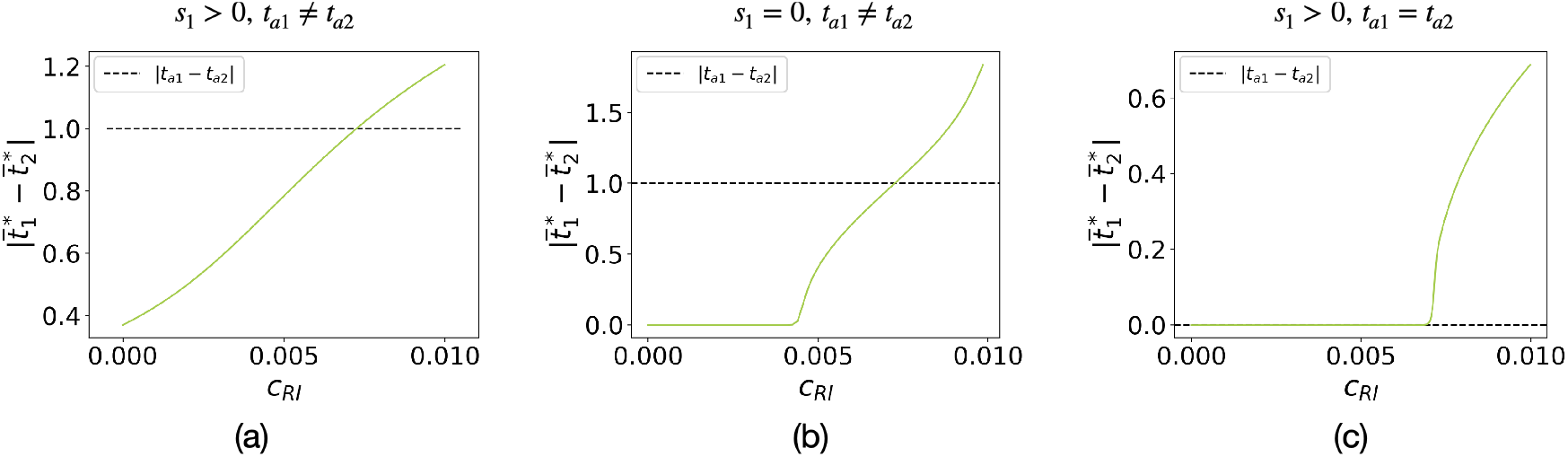
Influence of the cost of reproductive interference *c_RI_* on the phenotypic distances between the two species 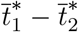 assuming strong female and predator discriminations. We assume (a) *s*_1_ = *s*_2_ = 0.005, *t*_*a*1_ = 0, *t*_*a*2_ = 1, (b) *s*_1_ = *s*_2_ = 0, *t*_*a*1_ = 0, *t*_*a*2_ = 1, and (c) *s*_1_ = *s*_2_ = 0.005, *t*_*a*1_ = *t*_*a*2_ = 1. We also assume: *G*_*t*_1__ = *G*_*p*_1__ = *G*_*t*_2__ = *G*_*p*_2__ = 0.01, *a*_1_ = *a*_2_ = *b* =1, *b* = 1, *d* = 0.02, *λ*_1_ = *λ*_2_ = 0.1, *n*_1_ = *n*_2_ = 10.

#### 6.5 Impact of predator discrimination without reproductive interference

**Figure A21:**
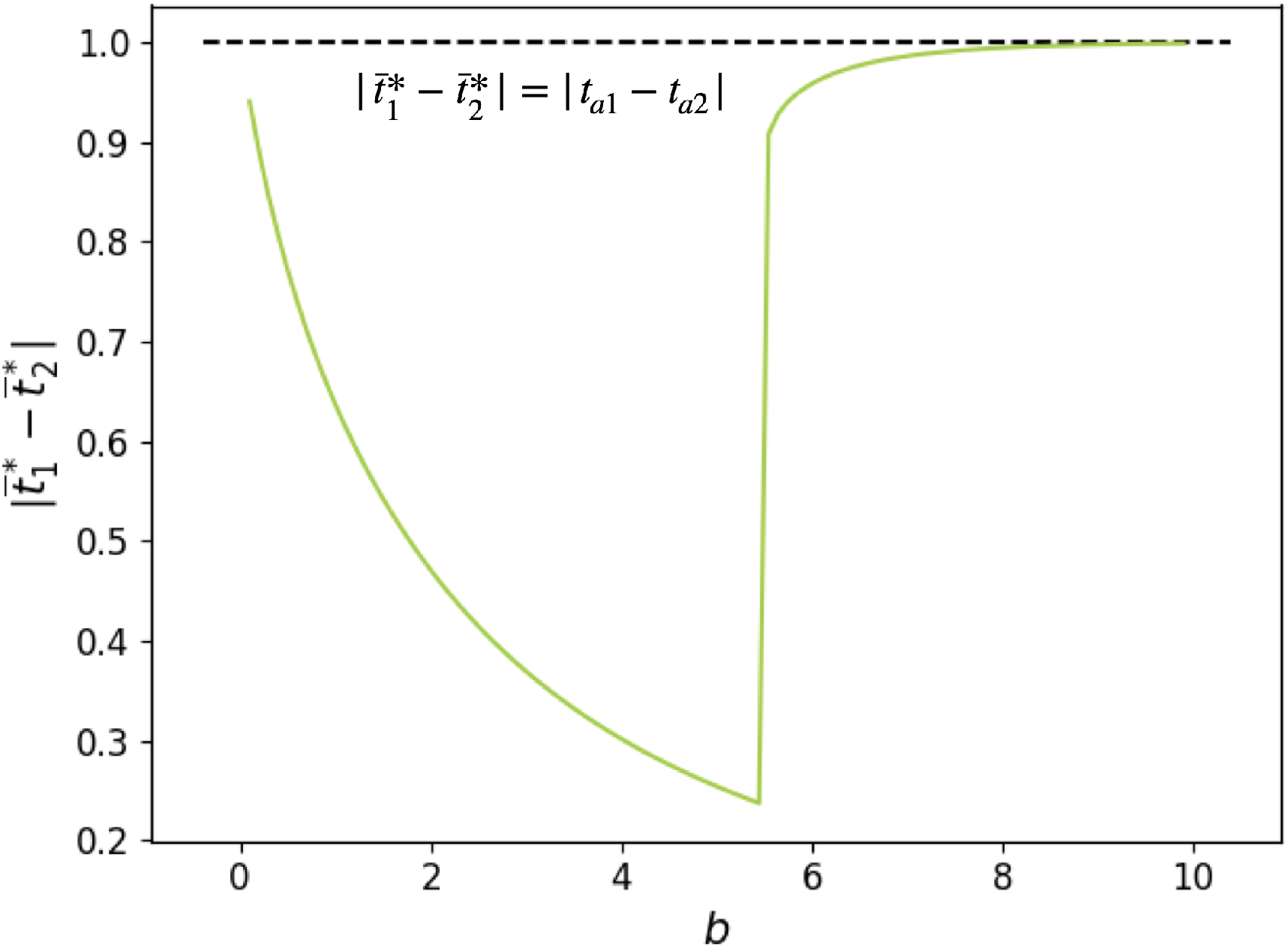
Influence of predator discrimination *b* on the phenotypic distances between the two species 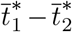 assuming no reproductive interference (*c_RI_* = 0). We assume: *G*_*t*_1__ = *G*_*p*_1__ = *G*_*t*_2__ = *G*_*p*_2__ = 0.01, *a*_1_ = *a*_2_ = *b* =1, *b* =1, *d* = 0.02, *λ*_1_ = *λ*_2_ = 0.1, *n*_1_ = *n*_2_ = 10, *s*_1_ = *s*_2_ = 0.005, *t*_*a*1_ = 0, *t*_*a*2_ = 1.

##### 6.6 Impact of the relative defence level on traits co-evolution

**Figure A22:**
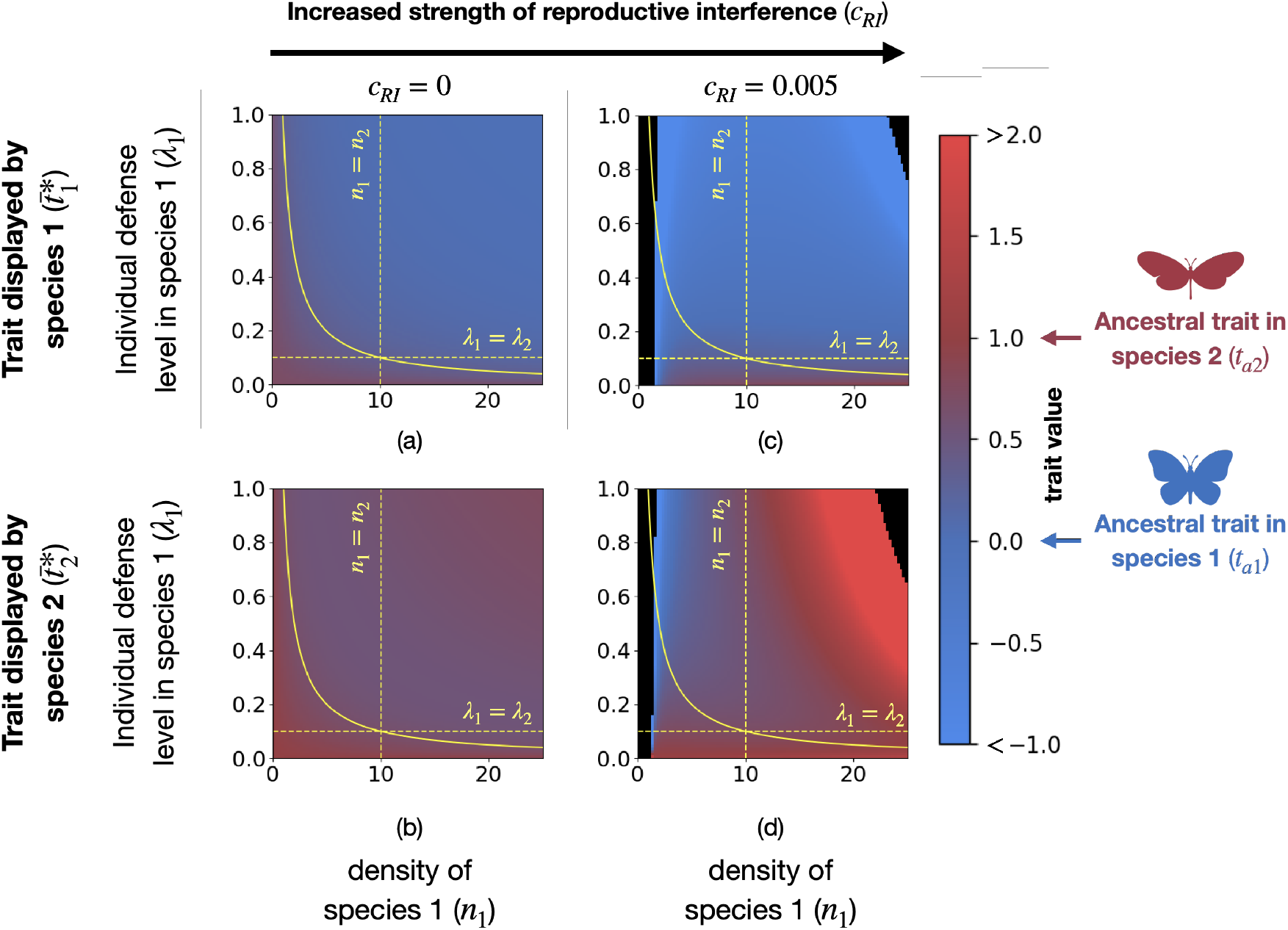
Influence of the density and of the individual defense level in species 1 (*n*_1_ and *λ*_1_) on the traits displayed in both species, for different strengths of reproductive interference (*c_RI_*). Plots report values obtained with the analytic approximation, assuming weak female and predator discriminations. Mean trait values can become very large (black zone). Trait values greater than 2 (resp. lower than −1) are shown in red (resp. blue). The yellow solid line shows the case where both species have the same level of defense (*λ*_1_*n*_1_ = *λ*_2_*n*_2_). Below (resp. above) this line species 1 has a lower (resp. higher) level of defense than species 2. Different values of strengths of reproductive interference are assumed: (a) and (b) *c_RI_* = 0, (c) and (d) *c_RI_* = 0.005. We assume: *G*_*t*_1__ = *G*_*p*_1__ = *G*_*t*_2__ = *G*_*p*_2__ = 0.01, *a*_1_ = *a*_2_ = 0.01, *d* = 0.05, *b* = 0.01, *λ*_2_ = 0.1, *n*_2_ = 10, *s*_1_ = *s*_2_ = 0.000005, *t*_*a*1_ = 0, *t*_*a*2_ = 1.

